# Systematic bias in surface area asymmetry measurements from automatic cortical parcellations

**DOI:** 10.1101/2025.03.25.645109

**Authors:** Yinuo Liu, Ja Young Choi, Tyler K. Perrachione

## Abstract

Anatomical asymmetry is a hallmark of the human brain and may reflect hemispheric differences in its functional organization. Widely used software like FreeSurfer can automate neuroanatomical measurements and facilitate studies of hemispheric asymmetry. However, patterns of surface area lateralization measured using FreeSurfer are curiously consistent across diverse samples. Here, we demonstrate systematic biases in these measurements obtained from the default processing pipeline. We compared surface area asymmetry measured from reconstructions of original brains vs. the same scans after flipping their left-right orientation. The default pipeline returned implausible asymmetry patterns between the original and flipped brains: Many structures were always left- or right-lateralized. Notably, these biases occur prominently in key speech and language regions. In contrast, manual labeling and curvature-based parcellations of key structures both yielded the expected reversals of left/right lateralization in flipped brains. We determined that these biases result from discrepancies in how regional labels are defined in the left vs. right hemisphere in the default cortical parcellation atlases. These biases are carried into individual parcellations because the FreeSurfer parcellation algorithm prioritizes vertex correspondence to the template atlas relative to individual neuroanatomical variation. We further demonstrate several straightforward, bias-free approaches to measuring surface area asymmetry, including using symmetric registration templates and parcellation atlases, vertex-wise analyses, and within-subject curvature-based parcellations. These results highlight theoretical concerns about using only the default processing stream to make inferences about population-level brain asymmetry and underscore the need for validating bias-free neuroanatomical measurements, particularly when studying regions where structural lateralization may underlie functional lateralization.

## Introduction

The left and right hemispheres of the human brain are similar but not symmetric in their structure. Hemispheric asymmetry is widespread and exists at both macrostructural and microstructural levels (Amunts, 2010). It is widely believed that structural asymmetries may underlie or reflect asymmetries in the brain’s functional organization (Toga & Thompson, 2003). For example, leftward asymmetry of superior temporal lobe structures has been associated with leftward asymmetry in language processing, based on measurements of both postmortem brain tissue (Galuske et al., 2000; Geschwind & Levitsky, 1968) and brain imaging data (Dorsaint-Pierre et al., 2006; Foundas et al., 1994; Josse et al., 2003; Ocklenburg et al., 2018; Shapleske et al., 1999; Sigalovsky et al., 2006; Tzourio, Nkanga-Ngila, & Mazoyer, 1998; Warrier et al., 2009). A large number of studies have also shown associations between altered brain structural asymmetry and a wide range of neurological, psychological, and psychiatric disorders, including autism (Floris et al., 2021), Alzheimer’s disease (Derflinger et al., 2011; Shi et al., 2009), dyslexia (Altarelli et al., 2014; Beaton, 1997; Robichon et al., 2000), and schizophrenia (Sommer et al., 2001).

At the macroanatomical level, human brain asymmetry is evident in hemispheric differences in the size, shape, and curvature of gyri and sulci (Renteria, 2012). Using magnetic resonance imaging (MRI), structural asymmetry can be measured *in vivo* by segmentation and parcellation of those structures (Van Essen et al., 2012). Widely used operationalizations of brain structure include *cortical surface area*, which measures the spatial extent of anatomically circumscribed brain areas (Winkler et al., 2012; Krubitzer, 2007); *cortical thickness*, which reflects local variation in the microstructural or laminar properties of the cortex (Fischl & Dale, 2000); and *volume*, which is an estimation of the amount of brain tissue within a region or structure (e.g., Filipek et al., 1989).

To obtain anatomical measurements, early studies depended on manual segmentation of volumetric brain scans carried out by expert neuroanatomists (e.g., Crespo-Facorro et al., 1999; Ebeling et al., 1989; John et al., 2006; Robichon et al., 2000; Tzourio, Nkanga-Ngila, & Mazoyer, 1998). More recently, powerful software packages that provide automated surface-based parcellation and analysis of cortical structure have become widely available, including *Caret* (Van Essen, 2012), *MindBoggle* (Klein et al., 2017), and *SUMA* (Saad et al., 2004; Saad & Reynolds, 2012). Perhaps the most widely used application for surface-based structural analyses is *FreeSurfer* (Fischl, 2012), which includes algorithms for volumetric segmentation, surface reconstruction, cortical parcellation, inter-subject alignment, and statistical analyses. Using these algorithms, researchers can easily obtain automatic measurements of myriad anatomical structures in individual brains, and these approaches have helped shed light on a wide array of important scientific and clinical questions (e.g., Bahar et al., 2023; Elliott et al., 2018; Kuperberg et al., 2003; Rimol et al., 2012; Silk et al., 2016; Tamnes et al., 2017; Wierenga et al., 2014; Williams et al., 2022; Yang et al., 2019). FreeSurfer’s automatic surface reconstruction is also integral to the preprocessing pipelines of many large-sample datasets, including the Adolescent Brain Cognitive Development (ABCD) Study and the Human Connectome Project (HCP) (Glasser et al., 2013; Hagler et al., 2019). In one such large-N study, researchers used the automated measurements from FreeSurfer to characterize putatively prototypical lateralization of key brain structures in a combined sample of over 17,000 healthy brains (Kong et al., 2018).

The automated analysis of structural lateralization offers a powerful analytical tool for large-N studies in many fields, including particularly the neurobiology of language, where the left-lateralization of language functions has been established for over 150 years (Broca, 1861; Wernicke, 1881). Such studies of structural asymmetries in speech and language areas have helped shed light on why leftward lateralization of language may exist and how abnormalities in lateralization may accompany language disorders (Altarelli et al., 2014; Elmer et al., 2013; Garnett et al., 2018; Greve et al., 2013; Marie et al., 2016; Meyer et al., 2014). Beyond language, automated structural parcellation has also been used to study endogenous patterns of brain lateralization (Kong et al., 2018; Kong et al., 2022) and how structural lateralization may be disrupted in various neurological or psychiatric disorders (Postema et al., 2019; Postema et al., 2021; Schijven et al., 2023; Sha, Schijven, & Francks, 2021).

However, comparing the surface area lateralization results across these studies raises an important concern about how these automated analyses are deployed: Different samples yield impossibly identical results when measuring hemispheric lateralization of surface area regardless of the sample size or population (**Fig. 1**). In this report, we examined whether there might exist hemispheric biases in the surface area measurements obtained from the default cortical reconstruction and parcellation stream in FreeSurfer that could explain why these measurements are so strikingly identical across diverse samples. Specifically, we compared cortical reconstructions and surface parcellations obtained from original volumetric MRI scans against those obtained using the same volumetric data with the left-right dimension reflected across the y-axis (i.e., “flipped” in the axial plane). We hypothesized that if the default reconstruction and parcellation pipeline were free from hemispheric bias, then the pattern of surface-area lateralization found in the flipped brains should be the *opposite* of that in the original brains (i.e., naturally left-lateralized structures should appear equally right-lateralized using the flipped data). We determined that systematic biases do, in fact, exist in the default cortical reconstruction and parcellation pipeline in FreeSurfer.

**Fig. 1:**
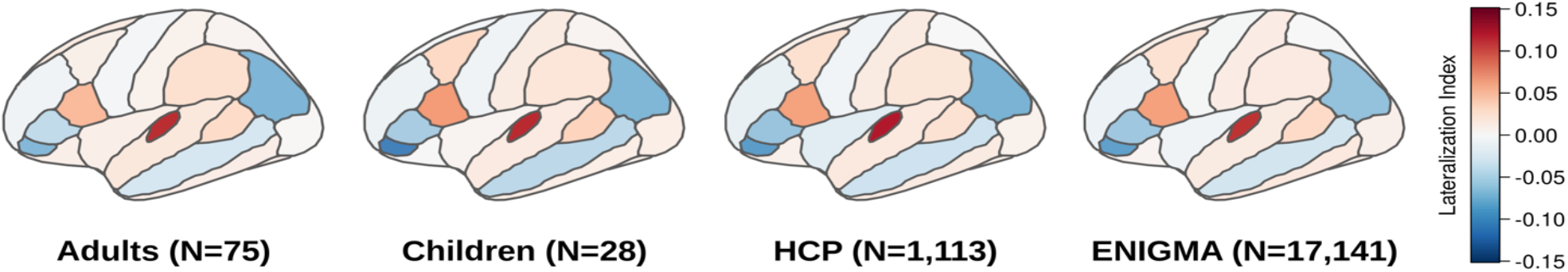
Strikingly identical patterns of cortical surface area asymmetry across different datasets obtained using the default *FreeSurfer* structural processing pipeline. The lateralization index was computed as in Eq. 1. Red hues represent leftward lateralization and blue hues represent rightward lateralization. The adult and child brains are sampled from Perrachione et al. (2016). HCP brains are from HCP S1200 dataset. ENIGMA brains are from Kong et al. (2018). We calculated the lateralization index for ENIGMA brains as a weighted average across datasets.

By systematically examining where, how, and why these biases occur, we determined that they result from discrepancies in how regional labels were defined relative to the curvature across the two hemispheres in the default parcellation atlases, especially the Desikan-Killiany atlas (better known as “*aparc*”) (Desikan et al., 2006). These biases in the parcellation atlases are carried forward into individual parcellation because FreeSurfer algorithms rely heavily on vertex alignment to the template surface and underweight individual curvature patterns. This leads to systematic bias that significantly overstates the true degree of left- or rightward surface area asymmetries in cortical regions. Critically, the magnitude of this bias is strongest in key structures that are relevant to speech and language, including especially inferior frontal gyrus (IFG) and transverse temporal gyrus (i.e., Heschl’s gyrus; HG). Finally, because this bias exists in the parcellation step of the default processing stream – and not the reconstruction or registration steps – we demonstrate how two straightforward changes (additions) to the default processing stream can be implemented to effectively ameliorate hemispheric biases and return more accurate values in cortical surface area measurements.

## Methods

### Participants

Structural brain images were obtained from *N* = 55 adult participants (38 female, 17 male; age 19-32 years, M = 22.6 years) recruited from the greater Boston University community. Participants reported tendency towards right-handedness (*n* = 43), left-handedness (*n* = 4) or ambidexterity (*n* = 8), as measured by the 10-item version of Edinburgh Handedness Inventory (Oldfield, 1971). This study was approved by the Institutional Review Board at Boston University and the Committee on the Use of Human Subjects as Experimental Subjects at the Massachusetts Institute of Technology. All participants provided written, informed consent and received monetary compensation for their time.

### MRI Data Acquisition

Using the Siemens Trio 3T scanner with a 32-channel phased array head coil at the Athinoula A. Martinos Imaging Center at the McGovern Institute for Brain Research at MIT, we obtained a whole-head, high-resolution T1-weighted (T1w), magnetization-prepared rapid gradient-echo (MPRAGE) anatomical volume (TR = 2,530ms, TE = [1.64, 3.50, 5.36, 7.22ms], TI = 1,400ms, flip angle = 7.0°, voxel resolution = 1.0mm isotropic, FOV = 256 × 256, 176 sagittal slices), as well as a matching T2-weighted (T2w) anatomical volume (TR = 3,200ms, TE = 454ms, voxel resolution = 1.0mm isotropic, FOV = 256 × 256, 176 sagittal slices). These data were collected as part of longer scanning sessions that involved unrelated diffusion and functional imaging not described here.

### MRI Data Analysis

Cortical reconstruction of the T1w anatomical images was performed using the default processing stream *recon-all* in FreeSurfer v6.0.0 (Dale et al., 1999). The T2w volumes were used to improve the estimation of the pial surface during cortical reconstruction. By default, the surfaces were registered to the built-in reference brain *fsaverage*, and different brain regions were automatically parcellated by built-in atlases, namely *aparc* (Desikan et al., 2006) and *aparc2009* (Destrieux et al., 2010) for cortical regions, and *aseg* (Fischl et al., 2002) for subcortical regions.

Using the same original T1w and T2w structural volumes above, we also created “flipped” copies using the FreeSurfer command *mri_convert*, in which the left-right directional encoding was reversed in the volume prior to submitting the scan to the FreeSurfer reconstruction pipeline. In this way, the original left hemisphere appeared in these flipped volumes as the “right” hemisphere, and vice versa. Hereafter, we will use *“left”* and *“right”* (in quotes) to refer to the apparent hemisphere in the flipped volume, and *left* and *right* (without quotes) to refer to the original (veridical) hemispheres. We then submitted the flipped T1w and T2w volumes to the same *recon-all* pipeline as above.

### Quantification of Bias

#### Lateralization Index and Bias Calculation

Using the cortical reconstructions and surface parcellations for both the original and flipped image volumes, we obtained the estimated cortical surface area and cortical thickness for each cortical region based on the default *aparc* atlas (Desikan et al., 2006) and *aparc2009* atlas (Destrieux et al., 2010), as well as the estimated tissue volume for each subcortical region based on *aseg* (Fischl et al., 2002). We analyzed the hemispheric lateralization of these anatomical measurements by computing the *lateralization index* (λ) for each region. This statistic is computed as the difference between the values obtained from the left and right (or “left” and “right” in the case of the flipped brains) hemispheres divided by their sum (Seghier, 2008):

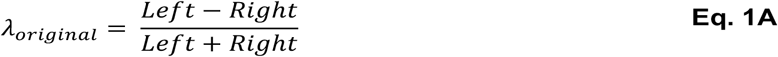

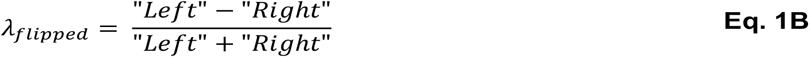

To quantify the degree of *lateralization bias* present in the values obtained from the default processing pipeline, we computed the differences between the lateralization index obtained from the original brains and the lateralization index obtained from the flipped brains for each cortical measurement and region per subject:

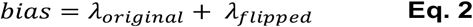

If no bias is present in the default parcellation, then the lateralization index of a region measured in a flipped brain should be the opposite of the lateralization measured in the original brain (e.g., having measured a leftward asymmetry of λ = 0.15 in a natural brain, we would expect an unbiased parcellation of that flipped brain to have a rightward asymmetry of λ = −0.15, such that unbiased λ_original_ + λ_flipped_ = 0). As such, positive bias values indicate systematic leftward asymmetries in the measurements obtained from the default automatic parcellations, and negative bias values indicate systematic rightward asymmetries. Because the lateralization indices are already scaled proportionately for every measurement (**Eq. 1A** and **Eq. 1B**), the magnitude of the bias values can be meaningfully compared across different regions, morphometric measures, and analytical variations.

### Hemisphere Classification

Previously, Hu et al. (2022) showed that training a support vector machine (SVM) based on surface area values obtained from the default processing stream in FreeSurfer could classify the left vs. right hemisphere of the human brain with a striking 99.4% accuracy. Following this work, we trained a linear SVM model using the HCP S1200 dataset from the Human Connectome Project (Van Essen et al., 2013), from which we used ∼70% (1,558 hemispheres) for training the model and the remaining 30% (668 hemispheres) for testing. We then used the trained model to classify the left vs. right hemispheres based on surface area values measured from our original and flipped brains.

When applied to the flipped brains in our study, the classifier should produce diametrically different results depending on whether it learned (1) something biologically real about surface area differences between the cerebral hemispheres or (2) any systematic left/right biases present in parcellations based on the default FreeSurfer processing pipeline. If the model learned to distinguish the true right and left hemispheres, then it should classify the “left” and “right” hemispheres of the flipped brains as the right and left hemispheres, respectively (i.e., apply the label reflecting the veridical hemisphere). However, if systematic hemispheric biases exist in the default surface area measurements, then we would expect the model to incorrectly classify the “left” and “right” hemispheres of flipped brains as the left and right hemispheres, respectively.

### Manual Parcellation

We wanted to compare surface area lateralization estimates from the automated pipeline against gold-standard manual segmentations of two prominent structures known to have reliable hemispheric asymmetries – HG and IFG. We manually labeled these structures on individual brains using the native-space pial and inflated surfaces for each hemisphere per participant. During manual labeling, the file names of the original and flipped brain surfaces were hidden using a random hash to avoid experimenter expectation bias while drawing surface labels.

The anterior boundary of HG was drawn along the first transverse sulcus and its posterior boundary along Heschl’s sulcus (Rademacher et al., 2001; Schneider et al., 2005; Warrier et al., 2009). The lateral extent of HG was truncated at the lateral surface of the superior temporal gyrus. For brains with HG duplications, the manual label included the sulcus intermedius, consistent with the labels in the default *aparc* atlas that do not distinguish gyri and sulci.

The posterior boundary of IFG was drawn at the inferior precentral sulcus and its superior boundary at the inferior frontal sulcus. The anterior boundary of IFG was either the lateral orbital sulcus or a vertical line between the inferior frontal sulcus and fronto-marginal sulcus, varying across participants. The inferior boundary was the Sylvian fissure and the circular sulcus of the insula. The subregions of IFG were divided by the rami of the Sylvian fissure. The horizontal ramus separates *pars orbitalis* from *pars triangularis*, while the ascending ramus separates *pars triangularis* from *pars opercularis* (Crespo-Facorro et al., 1999; Iordanova et al., 2023; John et al., 2006; Naidich, Valavanis, & Kubik, 1995; Yamasaki et al., 2010).

The surface area measurements of each manual label were obtained using the FreeSurfer command *mris_anatomical_stats*. The lateralization index and bias were calculated as **Eq. 1A, Eq. 1B**, and **Eq. 2** above. The reliability of manual labels was examined by using intra-class correlation coefficients (ICC) based on a two-way mixed effects model assessing absolute agreement (Yaakub et al., 2020) between the surface area estimations from same two images (e.g., original left hemisphere and flipped “right” hemisphere). The reliability of manual labels was also compared to the reliability of measures obtained via both the default processing stream and its modifications (see below).

### Amelioration of Bias

#### Symmetric Registration

To obtain cortical parcellations in FreeSurfer’s default reconstruction pipeline, each surface is first registered to a spherical template. The atlas-based parcellation is then computed with respect to this alignment. By default, there are two different registration templates, one for each hemisphere. It is possible that hemispheric biases are introduced by hemisphere-specific surface registration templates. When parcellating the flipped brains, the veridical left hemisphere was registered to the right hemisphere template and vice versa; it has been suggested that such contralateral registrations could result in alignment errors that might affect regional surface area estimation (Greve et al., 2013).

To examine the extent to which hemispheric differences in the surface-based registration templates were a source of lateralization bias, we registered all surfaces to the symmetric registration template *fsaverage_sym*, which is available in FreeSurfer but is not part of the default processing pipeline (Greve et al., 2013). We then used the left or right *aparc* atlas to parcellate the respective hemisphere using *mris_ca_label* based on the symmetric registration. This procedure allowed us to avoid registration bias in the flipped brains and to disentangle any registration bias from parcellation bias.

We examined surface registration as a source of lateralization bias in two ways. First, we obtained the lateralization index and bias as above for the cortical *aparc* regions based on the symmetric registration and default hemisphere parcellation. Second, we performed a vertex-wise analysis that measures the local surface area at each vertex independent of any parcellation scheme. This was done by registering all the left and right surfaces to the left *fsaverage_sym* so that the areal maps were all sampled with a deterministic correspondence between each vertex across hemispheres. A Jacobian correction was included during registration to account for stretching or compression (Winkler et al., 2012). The lateralization index was computed per vertex for each participant and the resulting individual lateralization map was surface-smoothed with a 10-mm FWHM Gaussian filter before averaging across the group (Maingault et al., 2016). We did this analysis for both the original and flipped brains.

#### Single-Atlas Parcellation

If hemispheric biases exist in the parcellation atlas, then using the same atlas to parcellate both hemispheres should reduce or remove parcellation differences as a source of bias (Li et al., 2024). To isolate the parcellation bias, we started with the symmetric registration and then used only the left *aparc* atlas to parcellate both hemispheres using *mris_ca_label*. To ensure correspondence between vertex sampling in the right surfaces and the left *aparc* atlas, parcellation of right hemispheres was carried out via Xhemi in FreeSurfer. The *λ*_*original*_ is calculated based on the parcellated original left hemisphere and Xhemi left hemisphere (which is the original right hemisphere), and similarly for *λ*_*flipped*_.

To reduce the concern about applying a specific hemispheric parcellation atlas to the contralateral hemisphere, we repeated the parcellation approach using only the right atlas. This also allowed us to compare how the same surface was parcellated differently by the left and right atlases. Since the surface’s curvature remains the same and the registration is symmetric, any difference in the parcellation can only be introduced by the parcellation atlas.

#### Individual Curvature-Based Parcellation

In contrast to the cortical parcellation in FreeSurfer, which is determined based on registration to a template surface, an alternative parcellation approach involves analysis of local curvature on native-space surfaces. Such an individual curvature-based labeling scheme is available for HG using the *Toolbox for the Automated Segmentation of Heschl’s gyrus* (TASH) (Dalboni da Rocha et al., 2020). We applied the TASH labeling scheme to both the original and flipped brains and obtained the lateralization index and bias for this parcellation of HG as above.

The first step in the TASH processing pipeline is to select candidate ROIs based on the *aparc2009* parcellations. Notably, the three regions defining the ‘raw auditory complex’ (HG, transverse temporal sulcus (HS), and planum temporale (PT)) are all affected by leftward bias in the *aparc2009* atlas (**Fig. S3**). Therefore, when refining the gyri in the selected ROIs, it is possible that a portion of HG in the right hemisphere is excluded from the candidate region, leading to systematically smaller HG on the right. To test if the curvature-based TASH parcellation also inherited systematic bias arising from the *aparc2009* atlas, we also applied the single-atlas-parcellation scheme on our symmetrically registered brains and ran the TASH parcellation based on the symmetric parcellation labels.

## Results

### Demonstration of Systematic Bias

If the automatic parcellation is unbiased, then applying the default processing pipeline to the original and flipped brain images should produce exactly opposite lateralization indices and thus lateralization bias values of zero. We tested the significance of lateralization biases via one-sample *t*-tests against zero in each cortical region.

We found that the default automated processing pipeline did have a significant hemispheric bias for nearly all regions (**Fig. 2** and **Table S1**). In *aparc*-based parcellations, many regions, including notably many key regions for speech and language, showed a significant leftward (and “left”-ward) lateralization in both the original and flipped brain images. For example, surface area measurements of HG, IFG *pars opercularis*, and the bank of the superior temporal sulcus (bank STS) were significantly and implausibly left-lateralized in both the original and flipped brains. Similarly, some regions, such as IFG *pars orbitalis* and IFG *pars triangularis*, showed implausibly consistent rightward (and “right”-ward) asymmetry in both the original and flipped brains.

**Fig. 2:**
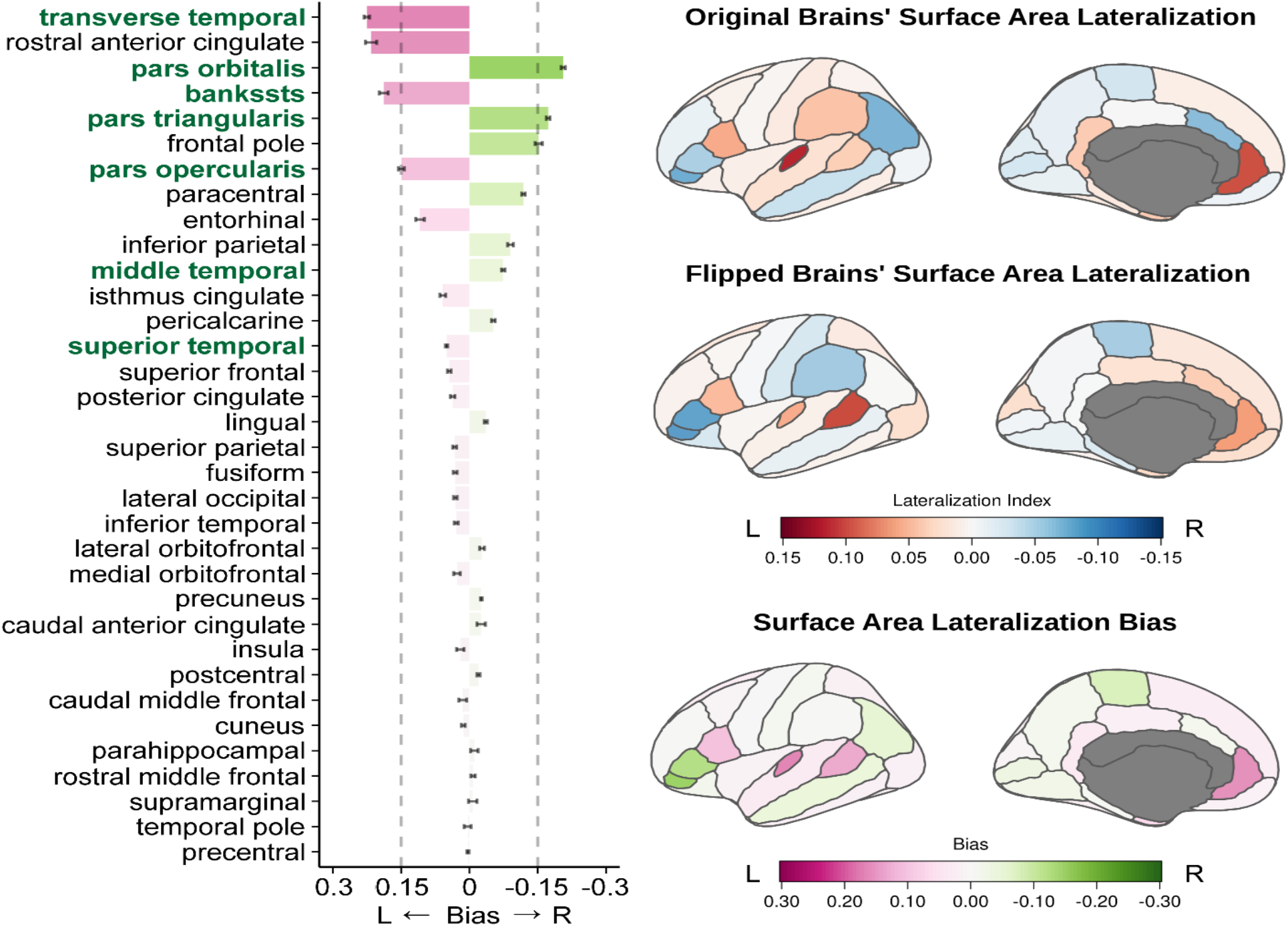
Systematic bias exists in the lateralization of cortical surface area measurements obtained using the default processing pipeline in FreeSurfer. Lateralization bias indicates implausibly consistent patterns of left- or rightward lateralization measured from reconstruction and parcellation of both original and flipped brain images. Key speech and language areas, indicated by green font, are especially affected by this bias. The dashed vertical lines anchor to the x-axis to help indicate the values.

While systematic lateralization biases were highly problematic in measurements of cortical surface area, we found that measurements of cortical thickness from *aparc* parcellations (**Fig. S1**) and subcortical volume from *aseg* segmentations (**Fig. S2**) exhibited much weaker biases. However, one-sample *t*-tests against zero nonetheless suggested that the bias was still significant for some regions (**Tables S2** and **S3**).

Cortical surface area lateralization biases exist not only in the default *aparc* atlas, but can also be found in the more granular *apac2009* atlas (**Fig. S3** and **Table S4**). Notably, the pattern of bias was not the same across these two atlases: Some areas, like HG, had similar leftward bias in both atlases, whereas others, like IFG *pars triangularis*, had opposite patterns of bias in the two atlases.

### Machine Learning Classification of Hemisphere

As shown previously (Hu et al., 2022), when we trained an SVM model to classify hemispheres as *left* or *right* based on surface area measurements from the HCP 1200 dataset, this classifier achieved an extremely (perhaps implausibly) high classification accuracy of 98.8% on an untrained testing sample of HCP brains. This model also achieved a strikingly high accuracy of 96.4% classifying the left vs. right hemisphere for the 55 original brains scans in this study. However, the model failed to correctly classify the veridical hemispheres of the flipped brains from our sample (i.e., correctly labeling the “left” hemisphere as the veridical right hemisphere), achieving an accuracy of only 20.9%. That is, in almost all cases, the classifier mislabeled the “left” hemisphere of the flipped brains as being the left hemisphere, despite the fact that it was actually the subject’s natural right hemisphere (and vice versa). We visualized the model coefficients from each region and noted that these values closely paralleled our measurement of systematic bias in each *aparc* region above (Pearson correlation: *r* = 0.850, *p* < 0.001) (**Fig. 3** and **Fig. S4**). These results indicate that the model did not learn the veridical hemispheric features, but instead learned the bias that exists between the left and right hemisphere versions of the parcellation.

**Fig. 3:**
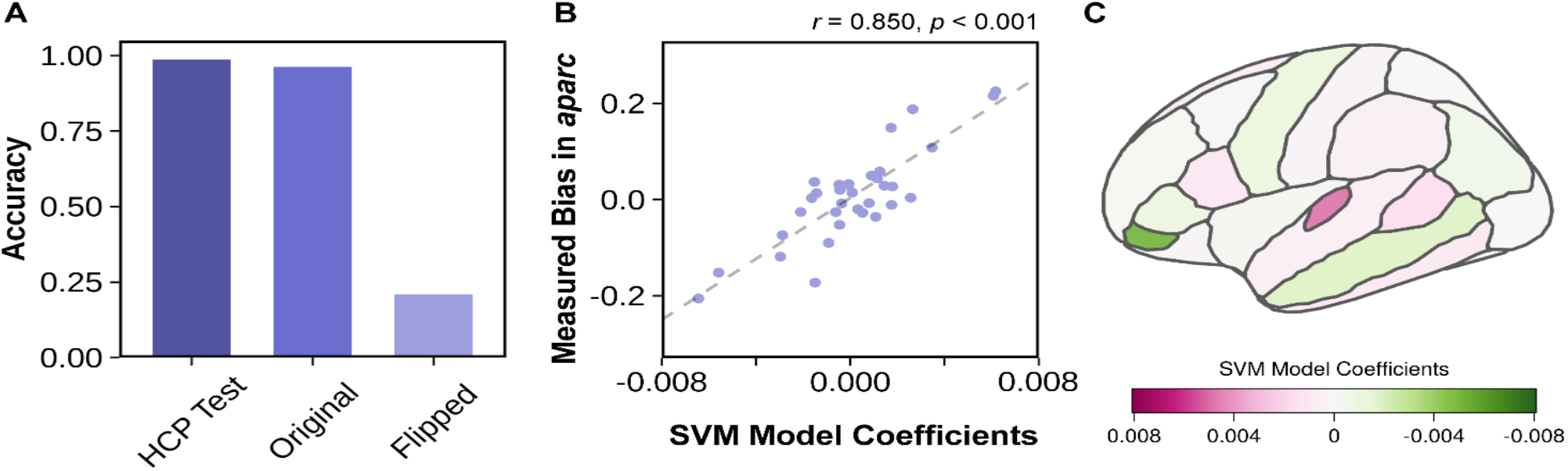
The linear SVM learned the systematic bias in the parcellation rather than real hemispheric differences. (A) The model achieved strikingly high accuracy when classifying the left vs. right hemisphere for the testing sample from the HCP as well as for the original brain scans in this study. However, it failed to correctly classify the veridical hemispheres of the flipped brains, indicating that the model did not learn the biologically real hemispheric features. (B)-(C) The SVM coefficients of each region closely correspond to the pattern of systematic bias we measured in the *aparc* atlas. The dashed line in (B) shows the regression fit. Compare the pattern of SVM coefficients in (C) to the regional bias in Figure 2.

### Comparison Between Automatic and Manual Segmentation

We manually labeled HG and IFG subregions for both the original and flipped brain images (**Fig. 4**). For HG, manual measurement of the original brains showed a slight left lateralization of this region (*t* = 1.731, *p* = 0.089), which is consistent with previous studies using manual segmentation (e.g., Kulynych et al., 1994, Marie et al., 2015). The lateralization index obtained using manual labeling was also much smaller than that obtained using FreeSurfer’s automatic pipeline (paired *t-*tests: original brains: *t* = 6.849, *p* < 0.001; flipped brains: *t* = 6.022, *p* < 0.001). Importantly, while manual measurement of the original brains found slight left-lateralization of HG, manual measurements of the flipped brains were slightly “right” lateralized (*t* = -1.535, *p* = 0.130), and there was no significant bias in the manual measurements (one-sample *t*-test vs. zero: *t* = 0.409, *p* = 0.684).

**Fig. 4:**
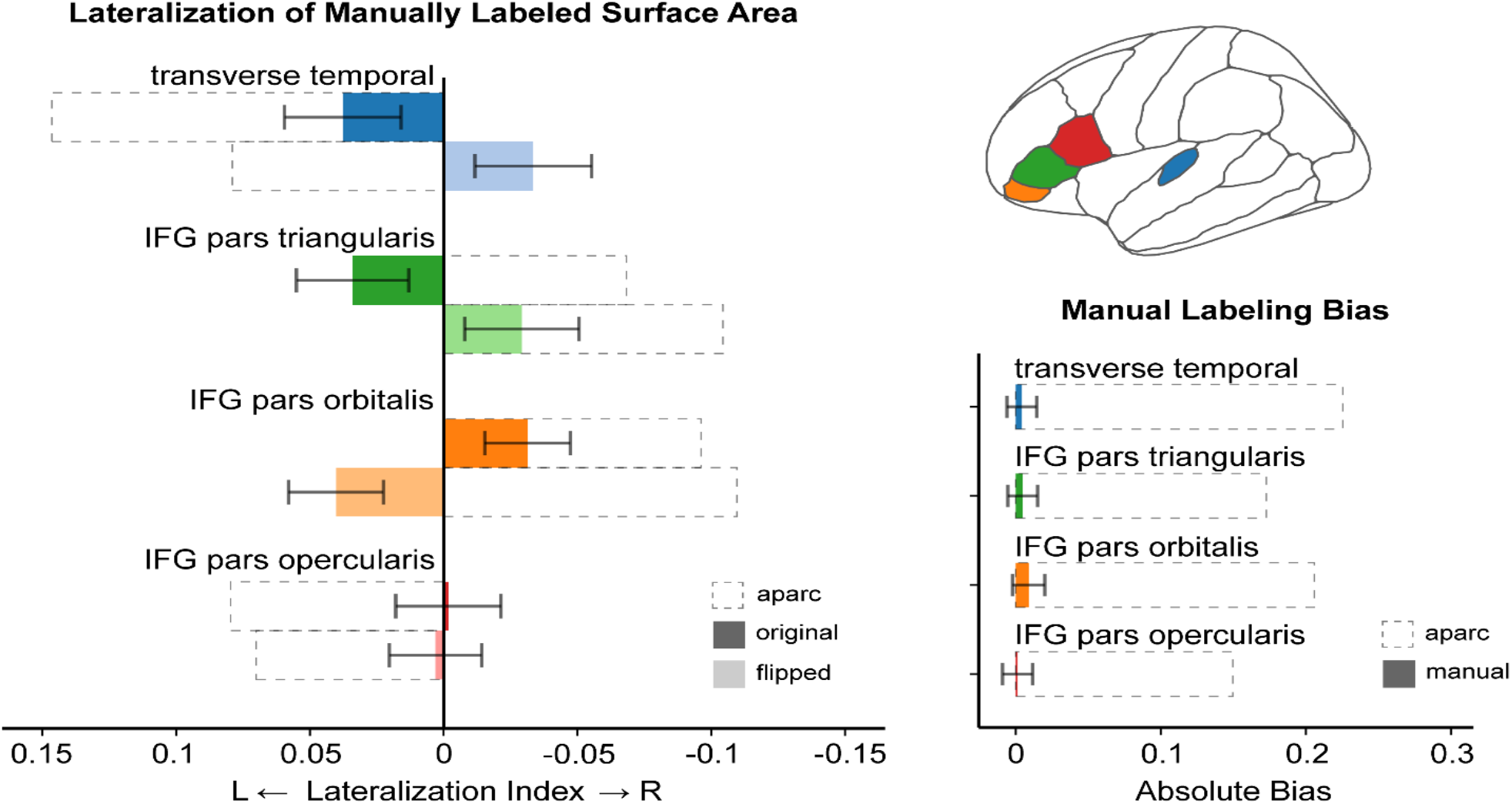
Opposite patterns of lateralization are found for original vs. flipped brains when manually labeling key regions based on established neuroanatomical landmarks. In contrast to automatic labeling based on the default processing stream, there was no hemispheric bias in regional labels that were drawn by hand. The magnitude of lateralization based on manual labeling was also smaller than that obtained using the default processing pipeline in FreeSurfer.

For manual segmentations of IFG in the original brain images, *pars orbitalis* showed a tendency towards rightward asymmetry (*t* = -1.963, *p* = 0.055); *pars triangularis* showed a slight left lateralization (*t* = 1.623, *p* = 0.110), which is contrary to the result from automatic parcellation; and *pars opercularis* showed no asymmetry (*t* = -0.090, *p* = 0.929), instead of the automatically measured leftward asymmetry. Similar to HG, the lateralization indices from manually labeled IFG subregions were much smaller than in the automatic parcellation (*pars opercularis*: *t* = 4.397, *p* < 0.001; *pars triangularis*: *t* = -4.718, *p* < 0.001; *pars orbitalis*: *t* = - 4.149, *p* < 0.001). As expected, the patterns of asymmetry were opposite in direction between the original and flipped brains (**Table S5**), and the hemispheric bias was effectively zero (*pars opercularis*: *t* = 0.122, *p* = 0.903; *pars triangularis*: *t* = 0.470, *p* = 0.640; *pars orbitalis*: *t* = 0.799, *p* = 0.428).

Using ICC, we determined that there was a high degree of consistency in the manual labeling for the veridical left and right hemispheres for each subject (i.e., the measurements made for the left hemisphere in the original brain and “right” hemisphere in the flipped brain). On the contrary, the automatic *aparc* labels showed poor reliability between the original and flipped hemispheres (**Table S6**), indicating that FreeSurfer produces different and inconsistent surface area measurements for a given region when the image was flipped.

### The Source of Systematic Bias

In the automatic parcellation pipeline, there are two steps where hemispheric biases may be introduced: registration and parcellation. We separated these two steps to identify the source of the bias.

We first registered all the surfaces to the symmetric template *fsaverage_sym* instead of the two separate, hemisphere-specific templates from *fsaverage* (Greve et al., 2013). If systematic biases exist in the registration step, then applying a symmetric registration should largely remove the observed bias with the default parcellation atlas. However, we found that the lateralization patterns in the original and flipped brains were still similar and the bias was significant (**Fig. 5**), indicating using a consistent surface template but still applying separate left and right parcellation atlases to the respective hemispheres did not reduce the bias (**Table S7**).

**Fig. 5:**
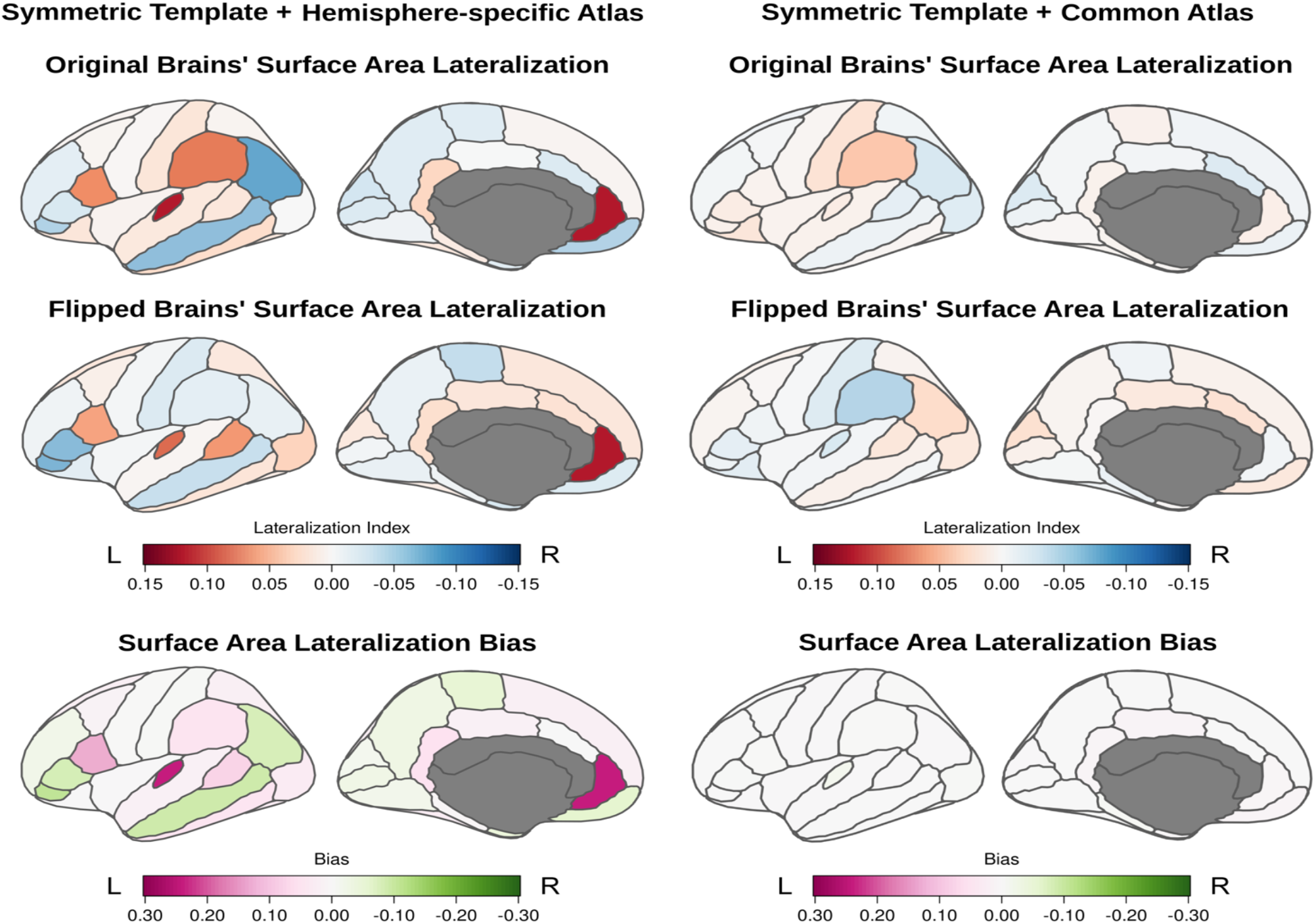
The bias exists in the parcellation step of the default processing stream and not the reconstruction or registration steps. The left column shows that using a symmetric registration template but retaining hemisphere-specific parcellations does not ameliorate the systematic biases in measurements of regional cortical surface area asymmetry. The right column shows that using a symmetric surface registration template plus parcellating both hemispheres with the same atlas (via X*hemi*) yields unbiased measures of regional cortical surface area asymmetry.

A vertex-wise analysis, which measures local surface area at each vertex without any parcellation, further confirmed that there was little hemispheric bias in the symmetric registration process: Left-lateralization of local surface area for corresponding vertices of the symmetric template in the original brains was paralleled by “right”-lateralization of the same vertices in the flipped brains, as seen prominently in regions like supramarginal gyrus and lateral occipital lobe (**Fig. 6**). These patterns are also consistent with parcellation, but not registration, as the principal source of bias.

**Fig. 6:**
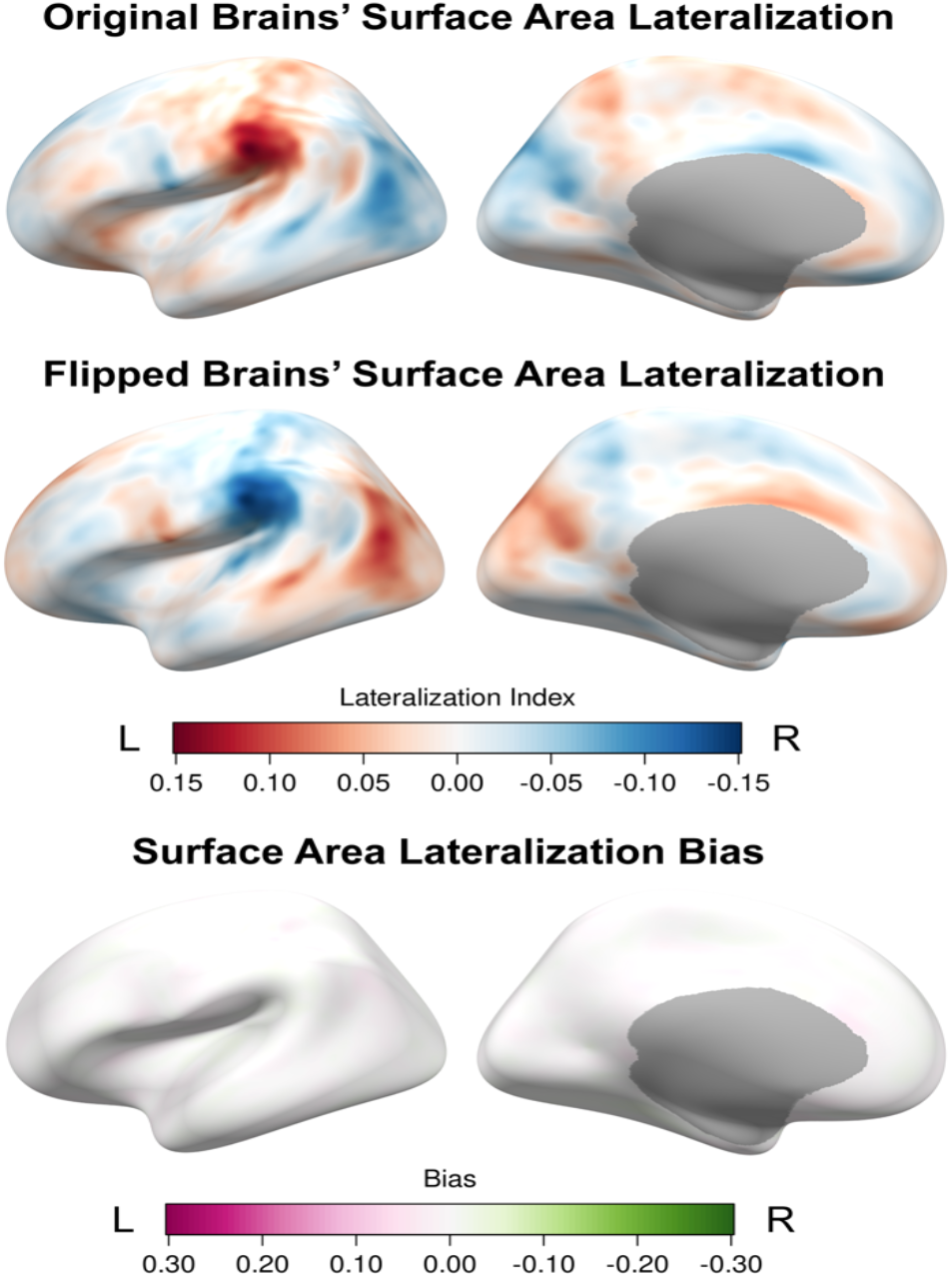
Vertex-wise lateralization analysis independent of any parcellation scheme did not show bias in the cortical surface area measurements. Individual left and right hemispheres were both registered to the left symmetric registration template (*lh*.*fsaverage_sym*) to ensure the correspondence between vertices in the left and right hemispheres.

If the systematic biases do exist in the parcellation atlas, then using the same atlas to parcellate both hemispheres should produce unbiased estimates of surface area lateralization. This was indeed what we found: Using a single atlas to parcellate both the left and right hemispheres significantly ameliorated the bias while preserving evidence of true hemispheric lateralization (**Fig. 5, Table S8** and **Table S9**). Using only either the left or right atlas does not affect the results (**Fig. S5** and **Table S10**). Additionally, the magnitude of surface area lateralization in both the original and flipped brains was much smaller than that obtained using the default processing pipeline. This magnitude, however, is much more similar to that obtained from manual labeling of the four regions we examined (HG, IFG *pars opercularis*, IFG *pars triangularis*, IFG *pars orbitalis*) (**Table S11**). These results confirmed the existence of systematic bias in the *aparc* parcellation atlas and that this bias exaggerates surface area lateralization.

We further examined why such biases occur in the parcellation atlases. By directly comparing the same surface (*fsaverage_sym*) parcellated using either the left or right *aparc* atlas, we identified significant discrepancies in how regional labels are defined relative to the curvature across the two hemispheres (**Fig. 7**). For example, IFG *pars opercularis* was separated from pars triangularis along the horizontal ramus in the right *aparc* atlas, but the boundary between these two regions is drawn much more anterior in the left *aparc* atlas. Correspondingly, this region was one with a significant leftward bias in the default *aparc* parcellation (**Fig. 2**), suggesting that the biases result from the discrepancies in regional boundaries’ definitions.

**Fig. 7:**
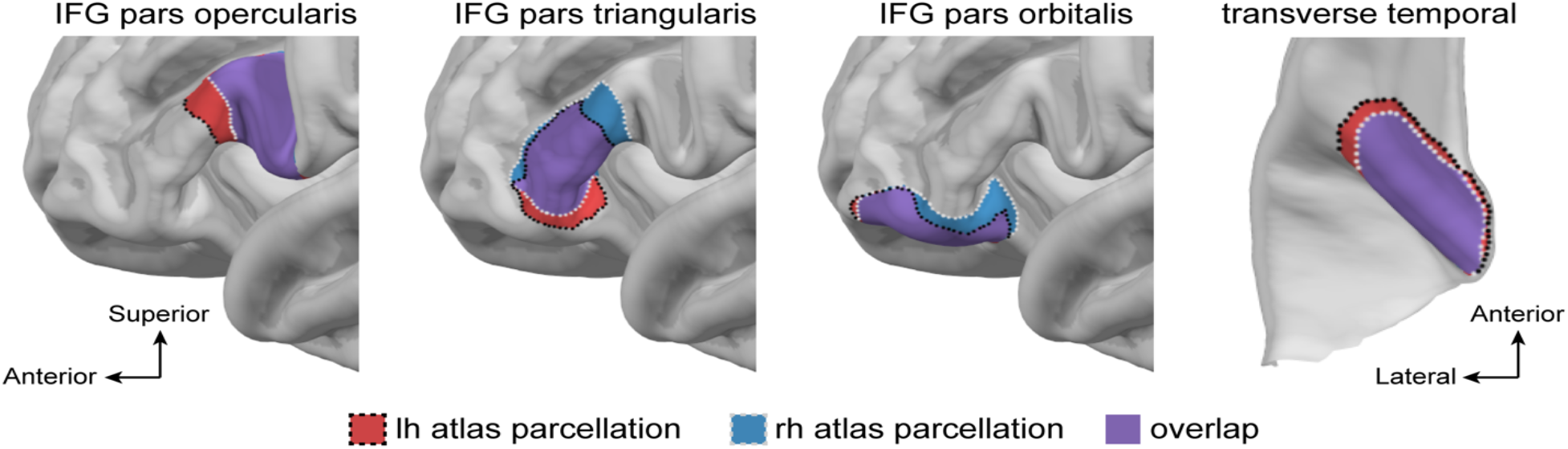
Discrepancies between the left and right *aparc* atlas in how regional labels are defined relative to the cortical curvature. The parcellation is shown on the *fsaverage_sym* surface, which has identical curvature across the two hemispheres, thus differences in the regional boundaries reflect only differences in the demarcation of these regions by hemisphere in the parcellation atlases. For example, the boundary of IFG *pars opercularis* was drawn substantially more anterior in the left *aparc* atlas, corresponding to the systematic leftward bias of this region.

### Native-Space Curvature-Based Parcellation

Although applying a single atlas to parcellate both hemispheres largely ameliorated the inherent biases in surface area lateralization measurements from the default processing stream, the automatic lateralization indices still differ slightly from those obtained via manual labeling (**Table S12**). These discrepancies arise largely from how regional labels are assigned by the FreeSurfer parcellation algorithm vs. human neuroanatomists: The FreeSurfer parcellation algorithm *mris_ca_label* prioritizes the prior probability of a vertex label when aligned to the surface template, rather than identifying the individual patterns of surface curvature that underlie classical labeling of brain areas by human observers. This heavily template-influenced labeling approach is evident when comparing “individually” labeled regions from native spaces in the *fsaverage* space. Individually assigned labels from the FreeSurfer algorithm result in a highly uniform overlap in the common space, whereas manual labels better highlight the extent of individual variability (**Fig. 8**). In fact, the individually drawn labels from FreeSurfer are only minimally different from labels obtained by simply transforming the *aparc* atlas from the surface template into native space using *mri_surf2surf* (Jaccard Index = 0.937). Because of the over-reliance on parcellation atlas, any asymmetry pattern in the atlas can be easily transferred into the individual parcellation (compare **Fig. 7** and **Fig. S6**). Accordingly, the systematic biases we measured were highly correlated with the lateralization indices of the *aparc* parcellation on the *fsaverage* template itself (*r* = 0.838, *p* < 0.001).

**Fig. 8:**
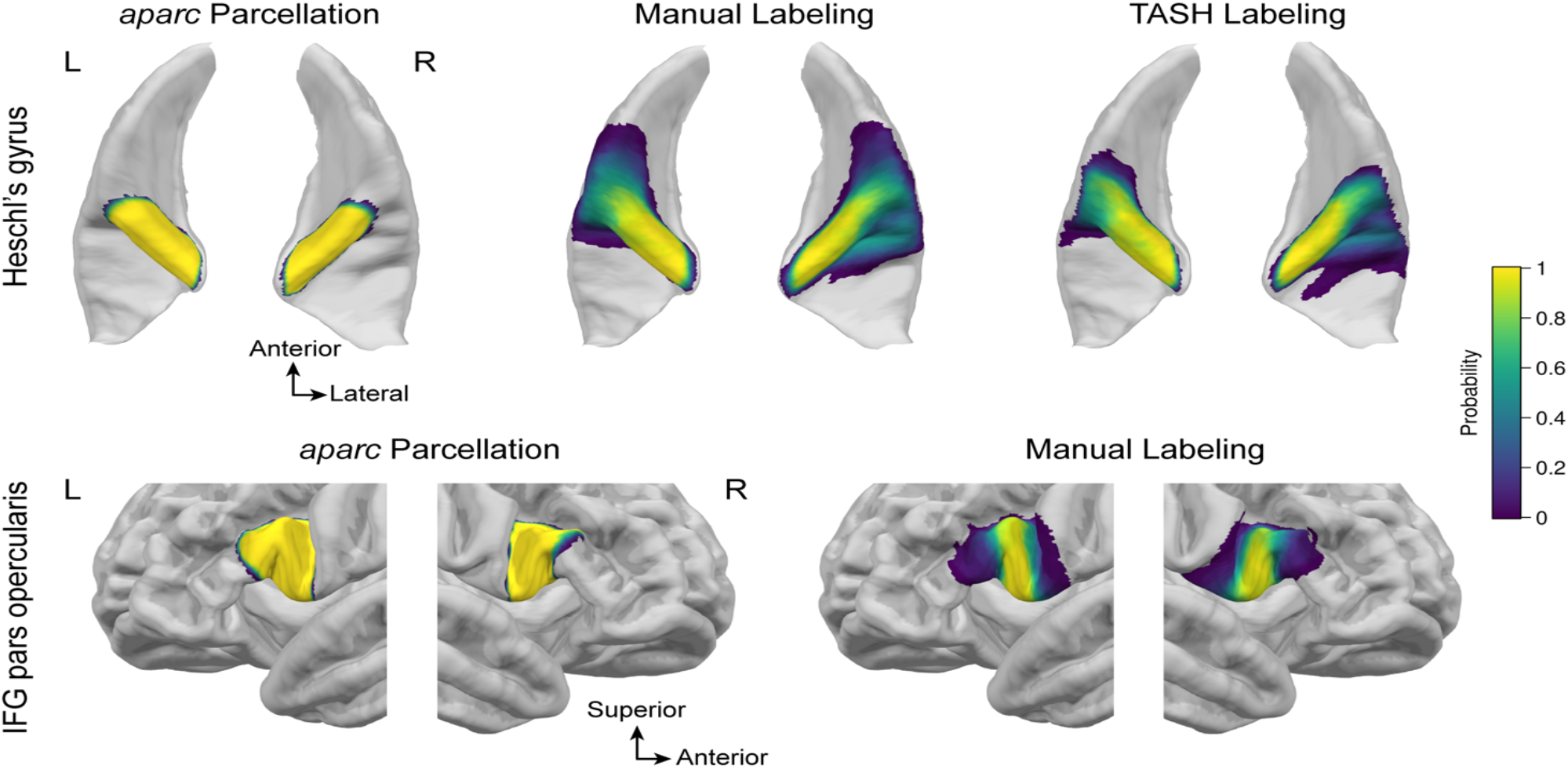
The default *aparc* parcellation is less sensitive to individual variation in the curvature and relies predominately on vertex-wise correspondence with the template. Mapping the regional labels from each individual’s brain obtained using the default processing pipeline in FreeSurfer back to *fsaverage* space reveals highly uniform overlap, demonstrating that even putatively individualized parcellations are in fact predominately derived from vertex-based alignment to the template rather than individual neuroanatomical landmarks. Any bias in the parcellation atlas will then be easily carried into individual parcellations. In contrast, overlaying the labels from each individual’s brain obtained via manual labeling or the bias-free curvature-based toolbox (TASH) in *fsaverage* space better reflects extent of individual variability in these structures.

A hybrid approach between vertex-based labeling and curvature-based labeling can be achieved using specialized schemes like those in the TASH parcellation for HG (Dalboni da Rocha et al., 2020). TASH starts with a circumscribed region based on an automatic parcellation, and then uses individual patterns of cortical curvature to identify HG. This approach is advantageous over labels based on alignment to a template, in that it can capture known patterns of variation in microanatomical structures, such as the various morphotypes of HG (Leonard et al., 1998). Although the final TASH labels include only the most anterior gyrus (including both branches of a common stem duplication), in the case of complete posterior duplication, a label for the second gyrus (HG2) is drawn separately and can be retained separately during the process. This approach captures the variability in HG morphology similarly to human labelers (**Fig. 8** and **Fig. S7**).

We used TASH to obtain HG labels for both the original and flipped brains based on individual patterns of curvature along the superior temporal plane. We calculated lateralization indices using the default output of TASH, which includes both branches of a common stem duplication but only the most anterior gyrus of a complete posterior duplication. There was significant leftward lateralization for the original brains (λ = 0.056, *t* = 2.784, *p* = 0.007). The magnitude was smaller than that obtained using default *aparc* parcellation (λ = 0.146; paired t-test: *t* = 5.072, *p* < 0.001) but similar to the estimates from manual labeling (λ = 0.038; paired *t*-test *t* = -0.796, *p* = 0.483). This leftward lateralization was mirrored by a tendency toward “right”-ward lateralization in the flipped brains, but this reversal was not significant (λ = -0.021, *t* = -0.616, *p* = 0.539). While largely reduced compared to bias in the default processing stream, some bias remained when labeling HG via TASH (one-sample t-test vs. zero: *t* = 1.877, *p* = 0.066, *d* = 0.253). The source of the remaining bias is related to the how TASH incorporates the *aparc2009* atlas in its initial processing steps. Modifying TASH to initialize from a symmetric parcellation helps reduce the bias (see **Supplementary materials**.)

## Discussion

In this study, we determined whether, where, and why systematic biases in cortical surface area measurements exist in the default FreeSurfer processing pipeline. Using reconstructions of brain volumes with the hemispheres “flipped” across the midline, we examined morphological measurements obtained from the automatic parcellations algorithms. We calculated lateralization indices for the original and flipped brain images and found that, for many regions, the direction and magnitude of surface area lateralization were in fact impossibly consistent between the original and flipped brains, demonstrating significant and widespread hemispheric biases in the surface area measurements obtained from the default processing stream.

Hemispheric bias from the automatic parcellation was also much larger for cortical surface area than for cortical thickness or subcortical volume. Manual measurements of surface area for several key structures revealed no such bias – the leftward lateralization found in the original brains was mirrored in rightward lateralization in the flipped brains when measured on the surface by hand. A linear SVM model trained to classify left vs. right hemispheres based on automatic regional surface area measurements was suspiciously accurate (>98%) for natural brains but mislabeled the vast majority (>80%) of flipped brains, further indicating biases in these measurements that do not reflect the underlying neuroanatomy. We then investigated whether the biases in the default processing pipeline arose due to hemispheric differences in the surface registration or parcellation steps. Surface area lateralization bias remained large regardless of whether the native space surfaces were registered to the default hemisphere-specific brain templates or to a symmetric template prior to parcellation. However, hemispheric bias was significantly reduced when using a single consistent atlas to parcellate both hemispheres, demonstrating that the source of these biases arise from the parcellation step when using the default parcellation atlas(es).

Detailed investigation of the left and right hemisphere versions of the default parcellation atlas *aparc* revealed discrepancies in how regional labels are defined relative to the curvature across the two hemispheres, especially in HG and IFG (Desikan et al., 2006; Fischl et al., 2004). For example, previous anatomical studies have described the boundary between IFG *pars orbitalis* and IFG *pars triangularis* as the anterior horizontal ramus (Crespo-Facorro et al., 1999; Foundas et al., 1998; Iordanova et al., 2023; Naidich et al., 1995). The division of IFG in the right hemisphere of the *aparc* atlas (Desikan et al., 2006) used by FreeSurfer is consistent with this definition. However, the division in the left hemisphere of the atlas is drawn much more anterior to this landmark, leading to a larger IFG *pars orbitalis* and a smaller IFG *pars triangularis* (**Fig. 7**) regardless of individual variation in the size of these structures. This pattern is consistent with our findings of a strong leftward bias for IFG *pars orbitalis* and rightward bias for IFG *pars triangularis* when estimating surface area using the default processing stream. Other regions show similar discrepancies in their demarcation vs. their nominal curvature-based landmarks between the two hemispheres. For example, the anterolateral extent of HG extends further into superior temporal gyrus in the left hemisphere version of the atlas (**Fig. 7**).

There can be two major sources for this inconsistency in anatomic boundaries between the left and right atlases. First, the *aparc* atlas is not strictly landmark-based, but instead the probabilistic result of training data. Early manual labeling studies have demonstrated the variability of gyral and sulcal patterns of IFG, where the *pars opercularis* can be divided into two gyri in the left hemisphere in some cases, and the anterior ramus separating the *pars triangularis* and *pars orbitalis* may not be apparent (Yamasaki et al., 2010). When averaging across the training set, the anatomical variations can lead to boundaries in the atlas that do not respect the patterns of curvature in either the template or in individual brains aligned to it. Another source might be related to the real lateralization in the training data. As indicated by our results, the regions carrying the bias are often speech and language regions. There are several studies using manual segmentation that show leftward lateralization for some language regions (e.g., Archila-Suerte et al., 2016; Cai et al., 2014; Curley et al., 2017; Gialluisi et al., 2017; Sheridan et al., 2022) using different measurements. Our manual labeling of HG also supports leftward surface area lateralization for this structure, with a lateralization index similar to that of earlier studies (Kulynych et al., 1994, Marie et al., 2015). As FreeSurfer’s default atlases were trained on manual segmentations of real brain images (e.g., the prior probability was obtained using the frequency of observing a certain label at a certain vertex), the asymmetry in the original anatomy can be carried into the atlases.

However, the problem is not that the atlases capture putatively real anatomical lateralization effects at the population level *per se*, but rather that the cortical labeling algorithm used by FreeSurfer carries these biases into individual parcellations irrespective of the actual underlying neuroanatomy of individual brains. Examining the labeling algorithm in FreeSurfer reveals how biases in the atlas get carried forward into individual parcellations: After registering individuals’ brains to a reference template based on an optimized alignment of curvature (Fischl et al., 1999), FreeSurfer assigns a regional level to each vertex in each hemisphere based on (1) the prior probability of each label occurring at each vertex, (2) the local spatial relationship between parcellation labels within some neighborhood of each vertex, and (3) the likelihood of observing the given surface geometry given the parcellation. This algorithm, implemented in *mris_ca_label*, prioritizes vertex correspondence to the template atlas much more than the pattern of individual curvature when assigning labels to individual brains. As we demonstrated, mapping the individually assigned labels from *aparc* parcellations back to the common space resulted in a highly concentrated probability map, whereas manual labels and curvature-based parcellations from TASH highlight the extent of individual variability in structures like HG (**Fig. 8** and **Fig. S7**). The individually drawn labels from FreeSurfer default reconstruction and parcellation pipeline via *mris_ca_label* were only minimally different from labels obtained by simply transforming the *aparc* atlas from the surface template into native space using *mri_surf2surf*. This can be problematic when individual curvature does not align well to the template (Duan et al., 2017). For example, FreeSurfer labels fail to capture individual variation in HG duplication (Marie et al., 2015), in contrast to approaches based strictly on individual curvature patterns, like manual labeling or TASH.

The presence of such systematic biases raise concerns about using the outputs of the default processing stream to characterize endogenous lateralization of the human brain (e.g., Kong et al., 2018) or to study group differences in lateralization (e.g., De Kovel et al., 2019) or structure-function correspondences between the hemispheres (e.g., Sha et al., 2021). In fact, the patterns of surface area lateralization found in large samples are effectively identical to the pattern of lateralization in the *aparc* atlas itself. A biased atlas can drive results, even from diverse samples, toward the same average “atlas asymmetry”, resulting in a much smaller difference between groups or a false negative result. This concern is particularly emphatic for investigations of regions related to speech, hearing, and language, which we found to be among those most strongly affected by the intrinsic biases in the default FreeSurfer atlases.

Given the popularity and relative ease of relying on FreeSurfer to investigate hemispheric asymmetry, we suggest that researchers exercise caution when drawing conclusions based on surface area values obtained from the default processing stream alone. Researchers may want to consider taking additional processing and quality control steps to insure the reliability of automated measures of cortical asymmetry, particularly for surface area. For example, following Greve et al. (2013) and Li et al. (2024), we show that using symmetric registration plus a symmetric parcellation (i.e., using a single atlas to parcellate both hemispheres) minimizes lateralization bias and yields lateralization magnitudes similar to gold-standard manual labeling. The symmetric registration template and the left or right parcellation atlases are available via FreeSurfer and easy to implement via additional steps after the default processing stream (see example code in the **Supplementary materials**). Although the accuracy of this approach is limited by the quality of alignment between the native and template surfaces, it does avoid introducing additional error from inconsistencies between hemisphere-specific atlases. Manual corrections to cortical labels can be made in the native space to correct for any errors between the atlas parcellation and native curvature – and the option to do so in contralateral registration via Xhemi may also help prevent introducing any experimenter expectancy effects.

Alternatively, researchers could rely on parcellations that are based primarily on landmarks in individual brain anatomy. We found that the curvature-based parcellation tool TASH segments HG more accurately than template-based parcellation in individual brains and can be implemented in a hemispheric bias-free way. There is need, however, for the development of similar toolboxes for other regions or ideally for whole-brain parcellation.

Finally, it is also important to note that although we only investigated the biases in *aparc, aparc2009*, and *aseg* atlases in this study, these biases could in principle exist in any atlas where left and right hemisphere parcellations were defined independently. Researchers using other software might also check for biases in their processing stream before making claims about endogenous lateralization, correlations with behavior, or differences between groups.

## Conclusion

We found that systematic hemispheric biases exist in cortical surface area measurements obtained from the default processing pipeline of FreeSurfer. We determined that these biases arise primarily from discrepancies in how the regional boundaries of homologous structures are defined in the default left vs. right *aparc* atlases. While these biases may reflect real population-level lateralization in the training data that was used to create these atlases, the way that FreeSurfer prioritizes generating single-subject atlas labels based on alignment to a template surface creates systematic error in how these parcellations are applied to individual brains. Critically, these biases are the largest in regions relevant to speech, language, and audition, where questions relating to individual or group differences in hemispheric lateralization are of considerable theoretical importance. Consequently, any biases present in the atlases end up not only in the statistics, but also in the theoretical conclusions we draw from them about functional neuroanatomy. To ameliorate these biases, researchers should conduct bias-free analyses based on symmetric parcellations, vertex-wise analyses, manual labeling, or within-subject curvature-based parcellations when making inferences about hemispheric lateralization of brain structures.

## Acknowledgments

We thank Terri Scott for data collection and Jason Tourville and Becky Belisle for helpful discussions. This work was supported by NIH grants HD096098 and DC014045 to TKP, and DC020208 to Zhenghan Qi. This project also made use of data provided by the Human Connectome Project, WU-Minn Consortium (Principal Investigators: David Van Essen and Kamil Ugurbil; 1U54MH091657) funded by the 16 NIH Institutes and Centers that support the NIH Blueprint for Neuroscience Research; and by the McDonnell Center for Systems Neuroscience at Washington University.

## Statements & Declarations

### Data Availability

The datasets analyzed during the current study are available from the corresponding author by reasonable request.

### Ethics Approval

This study was performed in line with the principles of the Declaration of Helsinki. Approval was granted by the Institutional Review Board of Boston University (Charles River Campus).

