## Supplementary Materials for "Systematic bias in surface area asymmetry measurements from automatic cortical parcellations"

**Supplementary Materials to accompany**
**Systematic bias in surface area asymmetry measurements from automatic cortical parcellations**

*Brain Structure and Function*

Yinuo Liu[1], Ja Young Choi[2], & Tyler K. Perrachione[1]

[1]Department of Speech, Language, and Hearing Sciences
Boston University
Boston, Massachusetts, USA

[2]Department of Communication Sciences and Disorders
Northwestern University
Evanston, Illinois, USA

**Contact**
Tyler Perrachione, PhD
635 Commonwealth Ave.
Boston, MA 02215

+1.617.358.7410


**TASH is partly affected by the bias in FreeSurfer atlas**

We used TASH to obtain HG labels for both the original and flipped brains based on individual patterns of curvature along the superior temporal plane. We found that hemispheric bias was largely reduced compared to the bias we observed in the default processing stream; however, some bias remained when labeling HG via TASH. We explored the algorithm further to examine the source of bias and how to ameliorate it.

Because the first step in TASH relies on candidate ROI selection based on *aparc2009* parcellations, which we also found to have systematic biases (**Table S4**), we therefore tested whether the observed bias in the curvature-based TASH parcellation was introduced at this step. We modified TASH to label HG based on applying a single-atlas-parcellation scheme to symmetrically registered brains. When parcellating both hemispheres using the right atlas only, there was significant leftward lateralization in the original brains ($\lambda = 0.059$, $t = 3.556$, $p < 0.001$), as well as significant "right"-ward lateralization in the flipped brains ($\lambda = -0.071$, $t = -4.018$, $p < 0.001$), and the bias was comparatively little ($t = -1.541$, $p = 0.129$).

Interestingly, when parcellating both hemispheres using the left atlas only, while the "right"-ward lateralization was still observed in the flipped brains ($\lambda = -0.063$, $t = -2.234$, $p = 0.030$), there was little to no lateralization in the original brains ($\lambda = -0.003$, $t = -0.123$, $p = 0.903$), leading to a large bias ($t = -2.403$, $p = 0.020$). By visually inspecting the candidate ROIs from each atlas, we noticed that left *aparc2009* atlas performed worse than the right atlas in demarcating the superior temporal plane by erroneously including too much of superior temporal gyrus (STG) within the candidate ROIs. Importantly, although the overall measurement of surface area was unbiased based only one single atlas, the defined boundary was slightly different across hemispheres. In the TASH pipeline, an "opening procedure" is adopted to remove these STG vertices (Dalboni da Rocha et al., 2020). In the original left hemisphere, the erroneously included STG was thin and can thus be removed successfully during this step. However, these formations included too many vertices and were not removed completely in the original right hemispheres, leading to a clearly erroneous parcellation of HG. Therefore, although TASH labeling is based on individual curvature and is unbiased itself, its accuracy is affected by the initial selection of candidate ROIs which relies on, and therefore is affected by bias in, the parcellation atlas.

**Example code to perform single-atlas parcellations**
To perform a single-atlas parcellation, you first obtain a symmetric registration for each subject, and then perform symmetric parcellation of those hemispheres using a single atlas (here, we show using the left atlas). Arguments that are hemisphere-specific have been bolded for clarity.

Create the Xhemi surfaces and the spherical registration lh.fsaverage_sym.sphere.reg:

```
# Original FreeSurfer reconstructions are in $SUBJECTS_DIR

# Create the subject's xhemi folders and populate them:
xhemireg --s $subject

# Register the left hemisphere to the left hemisphere of the symmetric template:
surfreg --s $subject --t fsaverage_sym --lh --no-annot

# Register the right hemisphere to the left hemisphere of the symmetric template:
surfreg --s $subject --t fsaverage_sym --xhemi --lh --no-annot
```

Perform the parcellation of both hemispheres using a single atlas:

```
# Parcellate the left hemisphere based on the left hemisphere atlas:
mris_ca_label \
       $subject \
       lh \
       $SUBJECTS_DIR/$subject/surf/lh.fsaverage_sym.sphere.reg \
       $FRESURFER_HOME/average/lh.DKaparc.atlas.acfb40.noaparc.i12.2016-08-02.gcs \
       $SUBJECTS_DIR/$subject/label/$subject.lh_atlas.sym.lh.annot

# Parcellate the right hemisphere based on the left hemisphere atlas:
mris_ca_label \
       $subject \
       rh \
       $SUBJECTS_DIR/$subject/xhemi/surf/lh.fsaverage_sym.sphere.reg \
       $FRESURFER_HOME/average/lh.DKaparc.atlas.acfb40.noaparc.i12.2016-08-02.gcs \
       $SUBJECTS_DIR/$subject/label/$subject.lh_atlas.sym.rh.annot
```

Note that to parcellate the right hemisphere, we used the Xhemi brains, such that calls to the "left" surface information refer to the veridical right hemisphere surface. Parcellation via the right hemisphere atlas can be achieved in a similar way, creating corresponding rh.fsaverage_sym.sphere.reg and substituting "rh" for all instances of "lh" in the calls to *mris_ca_label* above. Further clarification on the use of Xhemi can be found at https://surfer.nmr.mgh.harvard.edu/fswiki/Xhemi

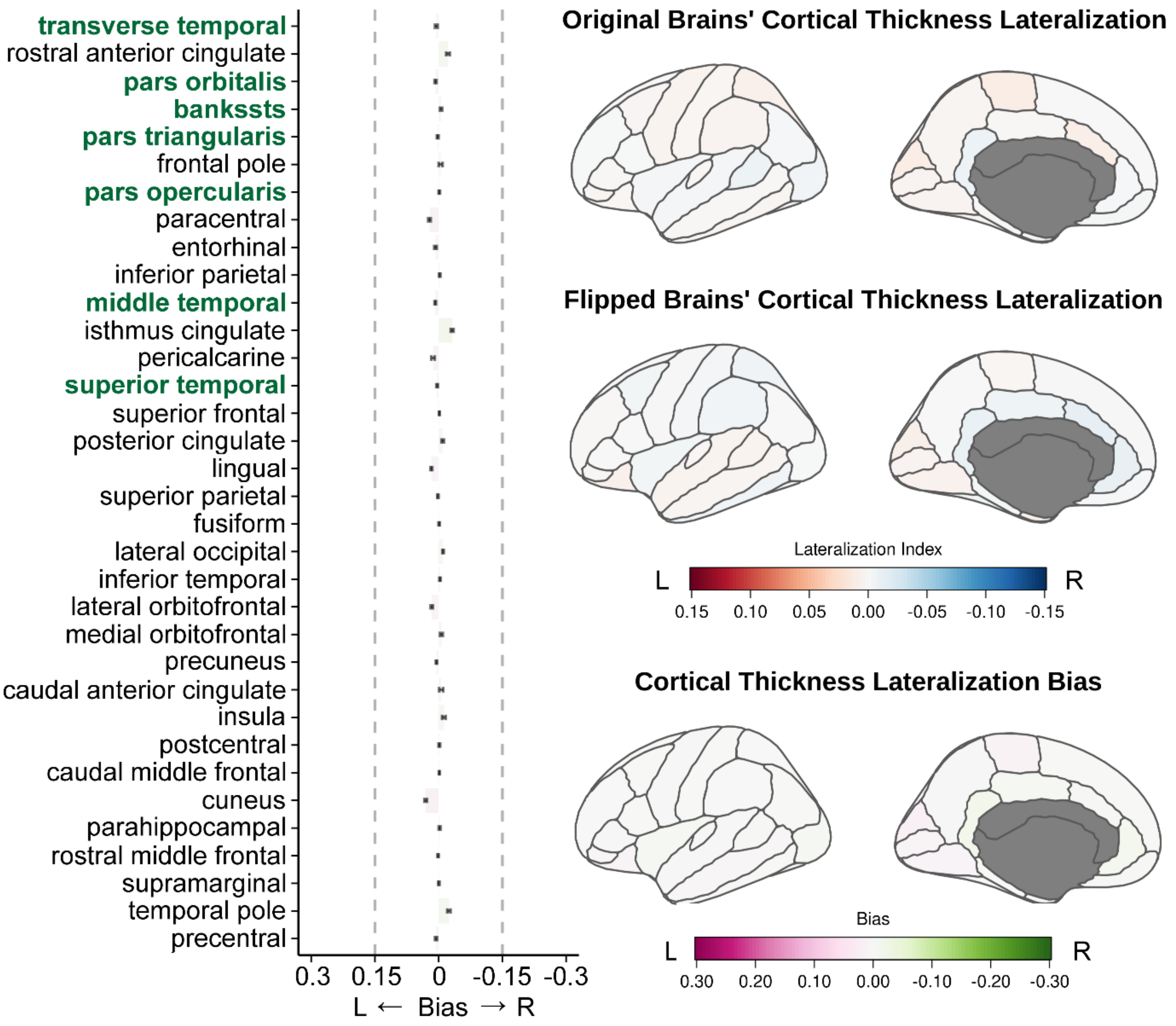

**Fig. S1: Comparatively little bias exists in measurements of cortical thickness lateralization obtained using the default processing pipeline in FreeSurfer.** Conventions as in Figure 2.

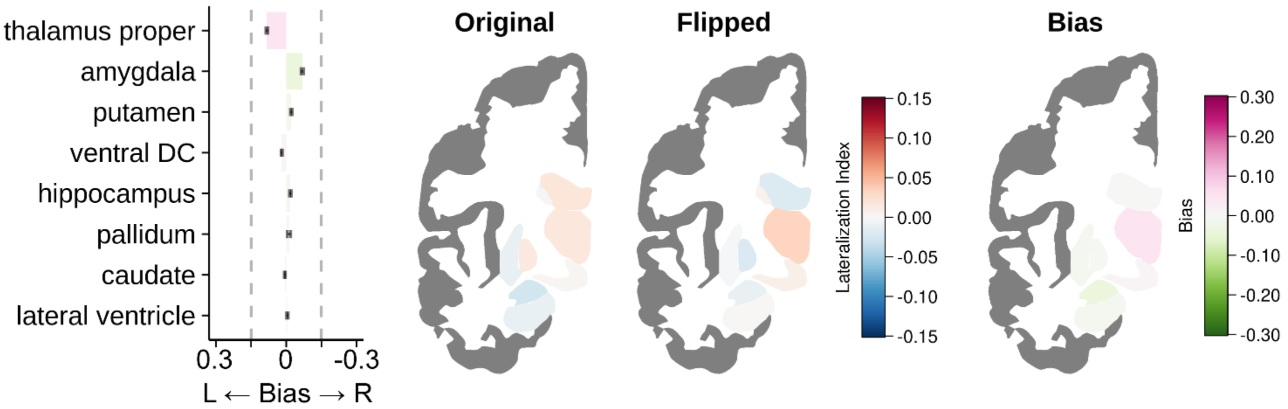

**Fig. S2: Smaller systematic bias also exists in some measurements of subcortical volume lateralization obtained using the default processing pipeline (*aseg*) in FreeSurfer.** Conventions as in Figure 2.

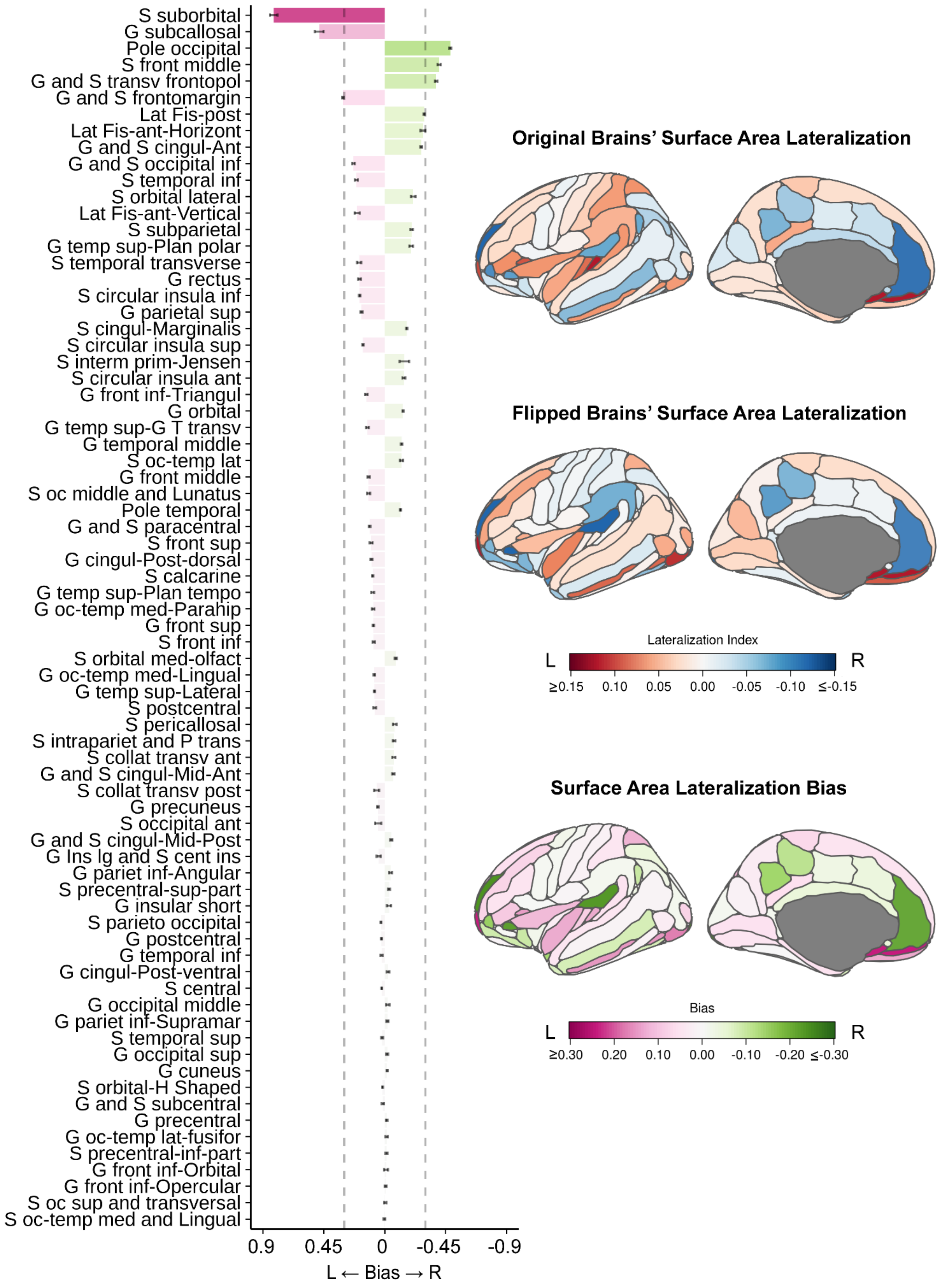

**Fig. S3: Systematic bias also exists in the lateralization of cortical surface area measurements obtained using *aparc2009* atlas in FreeSurfer but differs from the bias in *aparc* atlas.** The x-axis range of the bar figure differs from that in Figure 2, and the dashed lines indicate the range (±0.3) in Figure 2. For the brain figures, values outside the maximum or minimum of the range (±0.15 for lateralization index and ±0.3 for bias) are truncated for display.

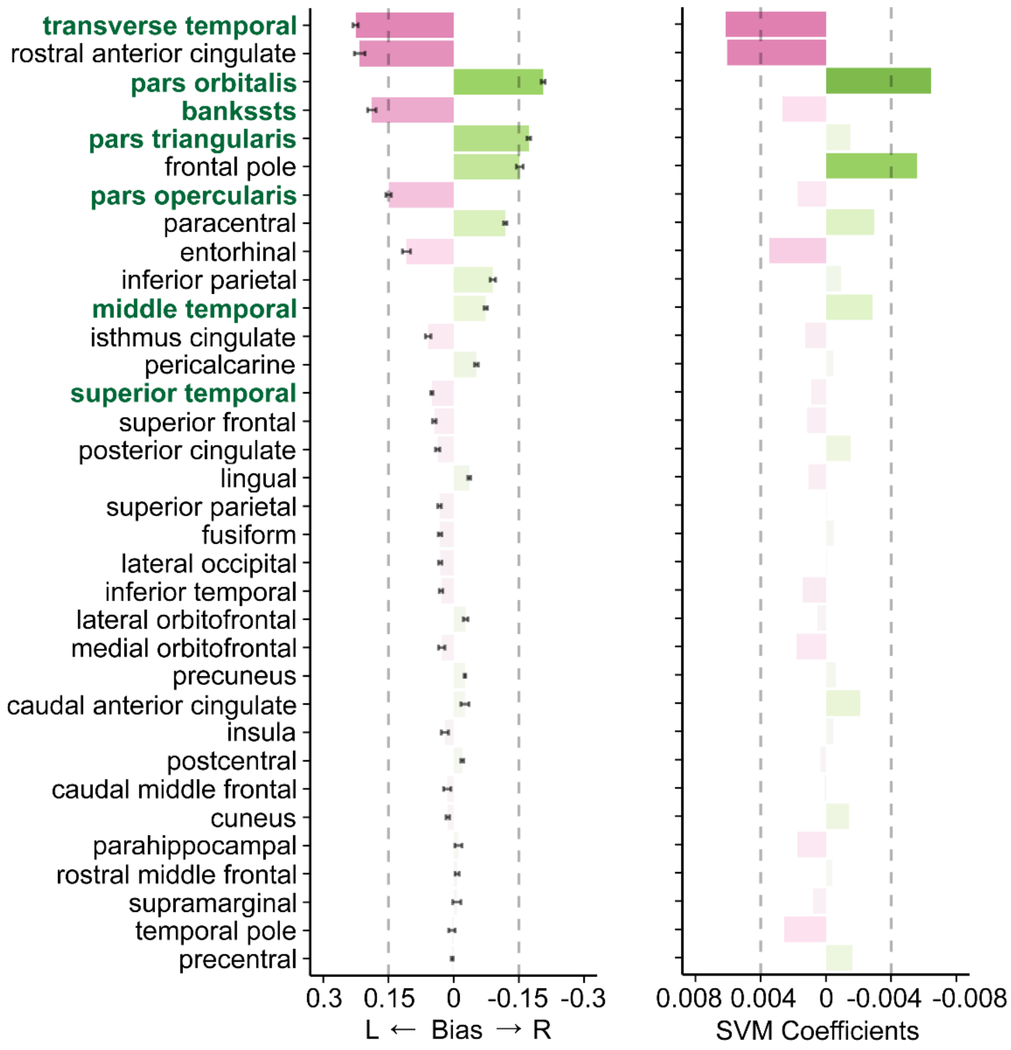

**Fig. S4: The SVM coefficients of each region show similar pattern of the systematic bias measured in *aparc* atlas.** This suggests that the linear SVM learned the systematic bias in the atlas parcellation rather than real hemispheric differences in the biology. The dashed vertical lines anchor to the x-axis to indicate the values.

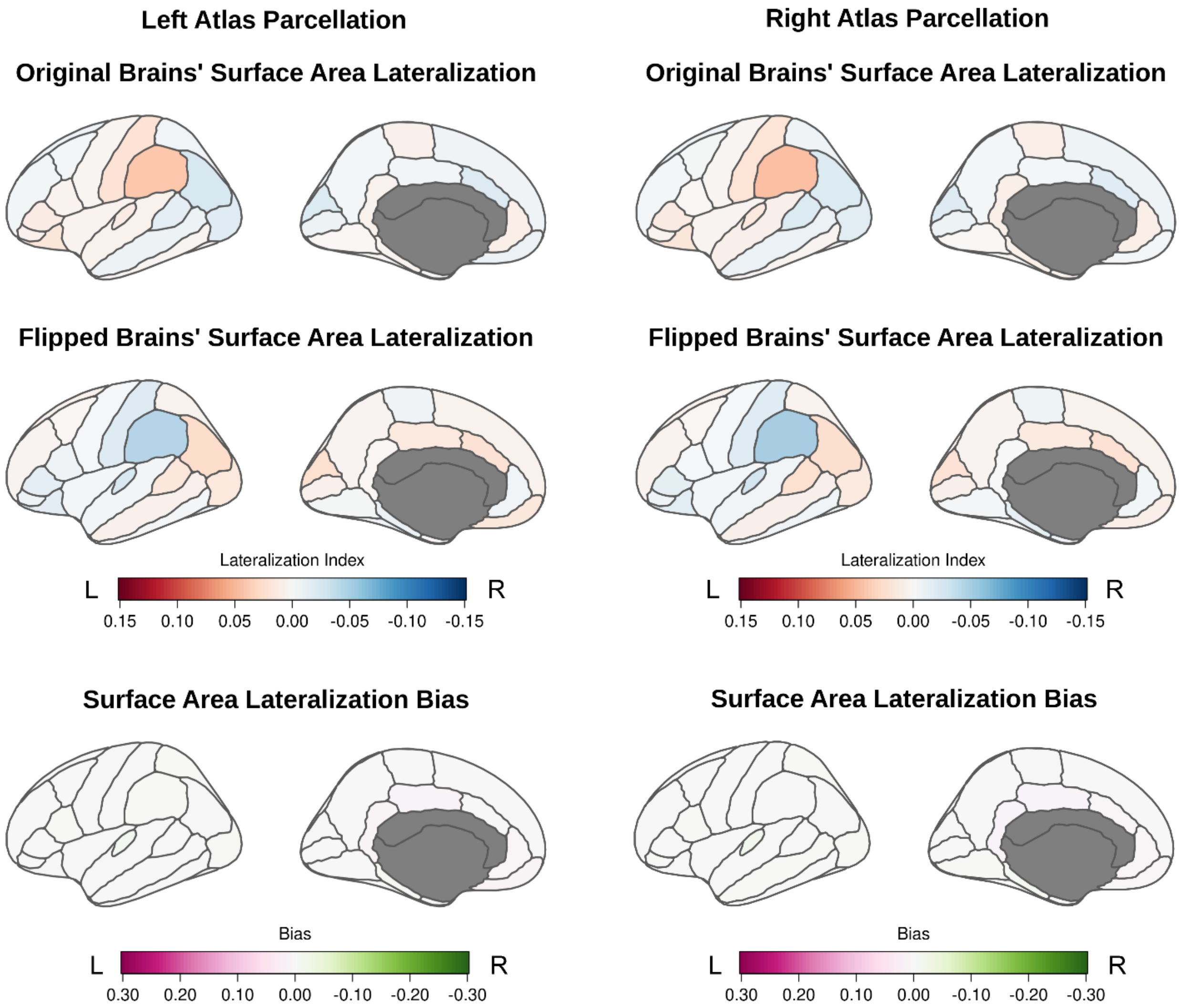

**Fig. S5: Using a symmetric surface registration template plus parcellating both hemispheres with the same atlas (via *xhemi*) yields unbiased measures of regional cortical surface area asymmetry**. The parcellation was done using either the left *aparc* atlas (left column) or the right *aparc* atlas (right column) after symmetric registration.

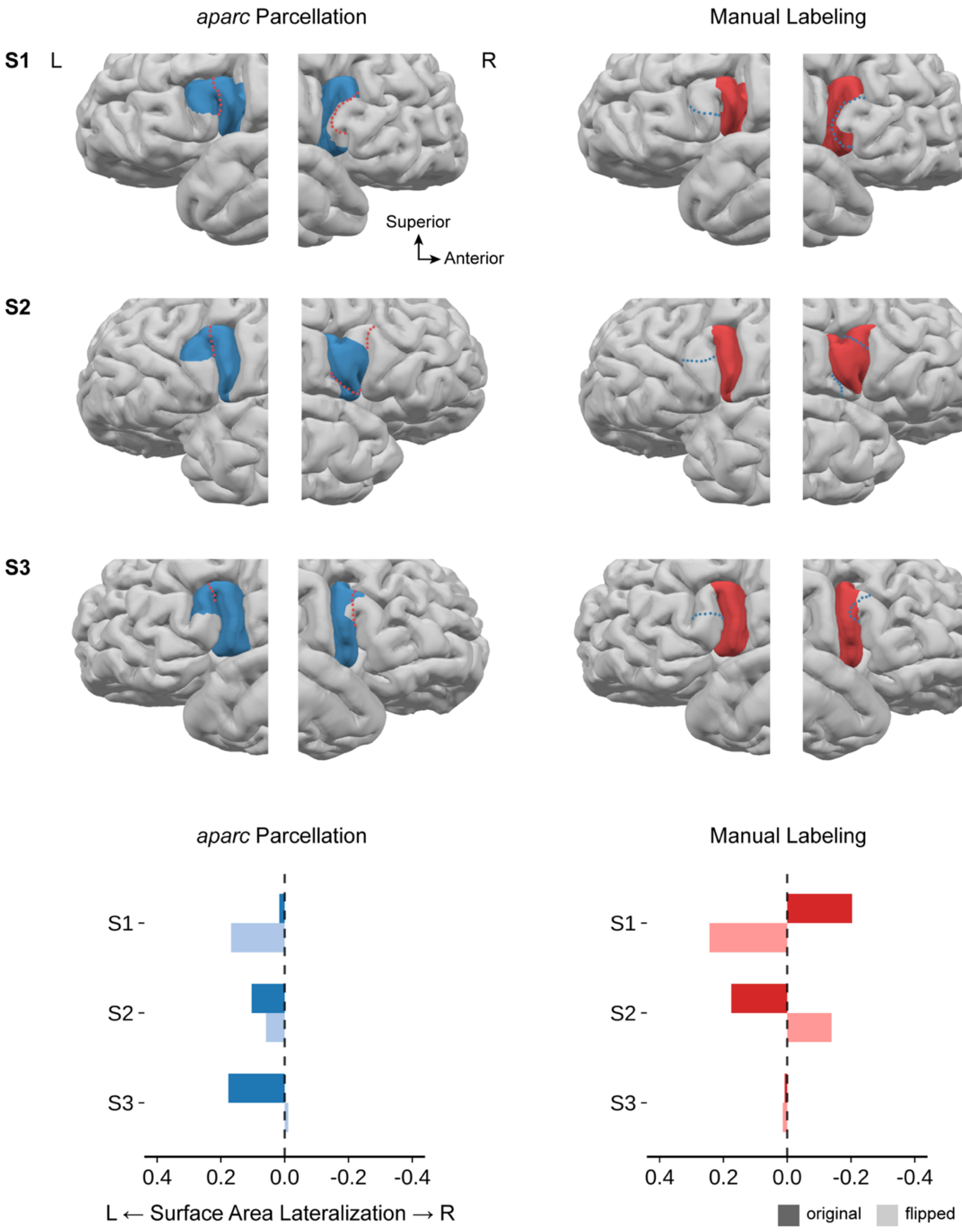

**Fig. S6: Examples of bias in IFG *pars opercularis* lateralization in individual brains labeled via FreeSurfer's default *aparc* parcellation vs. bias-free manual labeling.** Dashed lines show boundaries of corresponding labels by color. As shown in Fig. 7, the boundary of IFG *pars opercularis* was drawn substantially more anterior in the left *aparc* atlas. This bias is carried into individual labels by the *mris_ca_label* algorithm. On the contrary, manual labels showed the expected patterns of reversed lateralization with comparable magnitudes between the original vs. flipped brains.

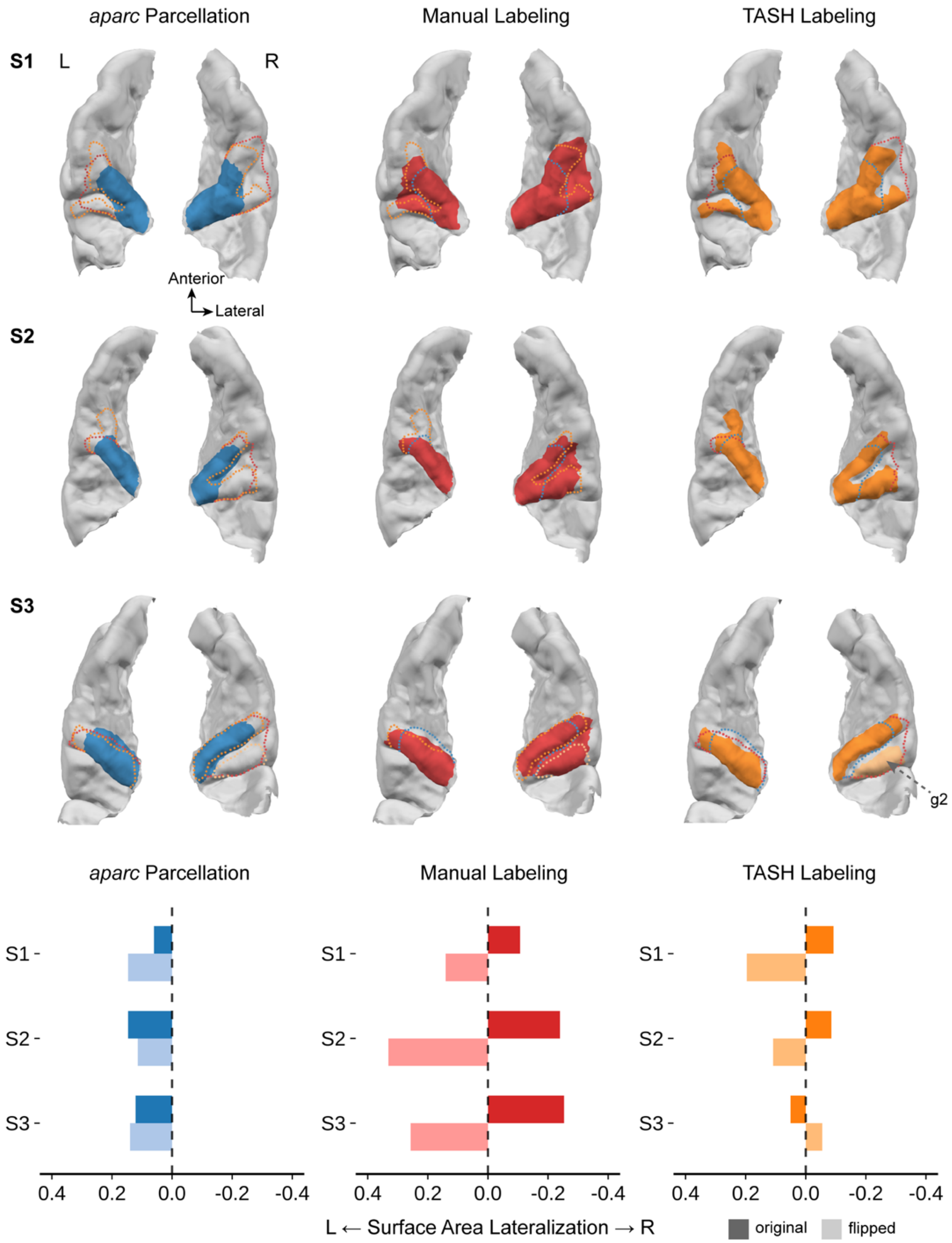

**Fig. S7: Examples of bias in HG lateralization in individual brains labeled via FreeSurfer's default *aparc* parcellation vs. bias-free manual and curvature-based toolbox (TASH) based labeling.** Both manual labels and TASH labels better capture the diverse configurations of HG and showed reverse patterns of lateralization in original vs. flipped brains, whereas *aparc* labels provided implausibly consistent leftward ("left"-ward) lateralization. Dashed lines show boundaries of corresponding labels by color. Note that TASH defines HG as gyri only, excluding Heschl's sulcus. In contrast to our manual labels, TASH deems only the most anterior gyrus as HG (Dalboni da Rocha et al., 2020), although it can separately segment the second gyrus of a complete posterior duplication. In such case (S3, right hemisphere), we plotted the selected anterior gyrus with orange and the identified but unselected posterior HG2 with a lighter shade. Only the selected gyrus was included in the lateralization calculation, resulting in a slight leftward lateralization of surface area in this case (vs. a strong rightward lateralization in the manual labels), but this difference is simply a matter of HG landmark definitions, which would be consistent between left and right (or left and "left") hemispheres, rather than parcellation issues as in *aparc* parcellations.

**Table S1:** Systematic bias in the *aparc* atlas when measuring cortical surface area lateralization using the default processing pipeline.

| Region | Mean Bias | *t* | *p* | Cohen's *d* |
|---|---|---|---|---|
| **bankssts** | 0.188 | 19.16 | <2.2e-16 | 2.58 |
| **caudal anterior cingulate** | -0.026 | -2.73 | 0.008 | 0.37 |
| **caudal middle frontal** | 0.015 | 1.71 | 0.094 | 0.23 |
| **cuneus** | 0.013 | 3.22 | 0.002 | 0.43 |
| **entorhinal** | 0.109 | 11.40 | 5.4e-16 | 1.54 |
| **frontal pole** | -0.152 | -18.80 | <2.2e-16 | 2.53 |
| **fusiform** | 0.031 | 7.41 | 9.0e-10 | 1.00 |
| **inferior parietal** | -0.090 | -13.83 | <2.2e-16 | 1.86 |
| **inferior temporal** | 0.029 | 6.82 | 8.0e-9 | 0.92 |
| **insula** | 0.020 | 2.41 | 0.019 | 0.32 |
| **isthmus cingulate** | 0.059 | 9.03 | 2.2e-12 | 1.22 |
| **lateral occipital** | 0.031 | 7.82 | 1.9e-10 | 1.05 |
| **lateral orbitofrontal** | -0.027 | -5.07 | 5.0e-6 | 0.68 |
| **lingual** | -0.036 | -10.44 | 1.4e-14 | 1.41 |
| **medial orbitofrontal** | 0.027 | 3.69 | 5.6e-4 | 0.50 |
| **middle temporal** | -0.074 | -19.08 | <2.2e-16 | 2.57 |
| **paracentral** | -0.119 | -30.22 | <2.2e-16 | 4.08 |
| **parahippocampal** | -0.011 | -1.30 | 0.200 | 0.18 |
| **pars opercularis** | 0.150 | 22.99 | <2.2e-16 | 3.10 |
| **pars orbitalis** | -0.206 | -44.39 | <2.2e-16 | 5.99 |
| **pars triangularis** | -0.173 | -40.51 | <2.2e-16 | 5.46 |
| **pericalcarine** | -0.052 | -12.28 | <2.2e-16 | 1.66 |
| **postcentral** | -0.020 | -5.68 | 5.6e-7 | 0.77 |
| **posterior cingulate** | 0.037 | 7.23 | 1.7e-9 | 0.97 |
| **precentral** | 0.003 | 1.43 | 0.157 | 0.19 |
| **precuneus** | -0.026 | -13.63 | <2.2e-16 | 1.84 |
| **rostral anterior cingulate** | 0.216 | 17.23 | <2.2e-16 | 2.32 |
| **rostral middle frontal** | -0.008 | -1.62 | 0.111 | 0.22 |
| **superior frontal** | 0.044 | 10.79 | 4.4e-15 | 1.45 |
| **superior parietal** | 0.032 | 7.59 | 4.5e-10 | 1.02 |
| **superior temporal** | 0.050 | 19.19 | <2.2e-16 | 2.59 |
| **supramarginal** | -0.007 | -0.76 | 0.453 | 0.10 |
| **temporal pole** | 0.004 | 0.52 | 0.603 | 0.07 |
| **transverse temporal** | 0.225 | 36.85 | <2.2e-16 | 4.97 |

Statistics reflect one-sample t-tests against zero.

**Table S2:** Systematic bias in the *aparc* atlas when measuring cortical thickness lateralization using the default processing pipeline.

| Region | Mean Bias | *t* | *p* | Cohen's *d* |
|---|---|---|---|---|
| **bankssts** | -0.006 | -2.05 | 0.045 | 0.28 |
| **caudal anterior cingulate** | -0.006 | -1.25 | 0.217 | 0.19 |
| **caudal middle frontal** | -0.001 | -1.24 | 0.218 | 0.17 |
| **cuneus** | 0.031 | 9.76 | 1.6e-13 | 1.32 |
| **entorhinal** | 0.007 | 2.56 | 0.013 | 0.35 |
| **frontal pole** | -0.004 | -0.99 | 0.327 | 0.13 |
| **fusiform** | -0.001 | -0.73 | 0.469 | 0.10 |
| **inferior parietal** | -0.002 | -1.66 | 0.103 | 0.22 |
| **inferior temporal** | -0.003 | -2.07 | 0.043 | 0.28 |
| **insula** | -0.012 | -2.91 | 0.005 | 0.39 |
| **isthmus cingulate** | -0.032 | -9.59 | 3.0e-13 | 1.29 |
| **lateral occipital** | -0.010 | -6.58 | 1.9e-8 | 0.89 |
| **lateral orbitofrontal** | 0.016 | 6.09 | 1.3e-7 | 0.82 |
| **lingual** | 0.172 | 8.37 | 2.5e-11 | 1.13 |
| **medial orbitofrontal** | -0.006 | -1.90 | 0.063 | 0.26 |
| **middle temporal** | 0.008 | 4.75 | 1.5e-5 | 0.64 |
| **paracentral** | -0.002 | -1.01 | 0.316 | 0.14 |
| **parahippocampal** | 0.022 | 7.26 | 1.5e-9 | 0.98 |
| **pars opercularis** | -0.001 | -0.89 | 0.378 | 0.12 |
| **pars orbitalis** | 0.007 | 3.17 | 0.003 | 0.43 |
| **pars triangularis** | 0.002 | 1.17 | 0.248 | 0.16 |
| **pericalcarine** | 0.013 | 3.23 | 0.002 | 0.44 |
| **postcentral** | -0.001 | -0.84 | 0.403 | 0.11 |
| **posterior cingulate** | -0.009 | -3.36 | 0.001 | 0.45 |
| **precentral** | 0.006 | 2.67 | 0.010 | 0.36 |
| **precuneus** | 0.005 | 3.98 | 2.1e-4 | 0.54 |
| **rostral anterior cingulate** | -0.022 | -5.26 | 2.4e-6 | 0.71 |
| **rostral middle frontal** | 0.001 | 1.05 | 0.299 | 0.14 |
| **superior frontal** | -0.002 | -1.70 | 0.096 | 0.23 |
| **superior parietal** | 0.002 | 1.25 | 0.216 | 0.17 |
| **superior temporal** | 0.004 | 1.95 | 0.056 | 0.26 |
| **supramarginal** | 0.000 | -0.28 | 0.780 | 0.04 |
| **temporal pole** | -0.024 | -6.20 | 8.2e-8 | 0.84 |
| **transverse temporal** | 0.006 | 1.72 | 0.091 | 0.23 |

Statistics reflect one-sample t-tests against zero.

**Table S3:** Systematic bias in the *aseg* atlas when measuring subcortical volume lateralization using the default processing pipeline.

| | **Mean Bias** | ***t*** | ***p*** | **Cohen's *d*** |
|---|---|---|---|---|
| **amygdala** | -0.069 | -12.73 | <2.2e-16 | 1.72 |
| **caudate** | 0.006 | 2.09 | 0.041 | 0.28 |
| **hippocampus** | -0.018 | -4.25 | 8.5e-5 | 0.57 |
| **lateral ventricle** | -0.005 | -1.33 | 0.188 | 0.18 |
| **pallidum** | -0.012 | -1.68 | 0.098 | 0.23 |
| **putamen** | -0.022 | -5.43 | 1.3e-6 | 0.73 |
| **thalamus proper** | 0.083 | 18.46 | <2.2e-16 | 2.49 |
| **ventral diencephalon** | 0.020 | 6.47 | 3.0e-8 | 0.87 |

**Table S4:** Systematic bias in *aparc2009* atlas when measuring cortical surface area lateralization using the default processing pipeline.

| | Mean Bias | *t* | *p* | Cohen's *d* |
|---|---|---|---|---|
| **G and S frontomargin** | 0.309 | 51.39 | <2.2e-16 | 6.93 |
| **G and S occipital inf** | 0.231 | 24.37 | <2.2e-16 | 3.29 |
| **G and S paracentral** | 0.112 | 17.10 | <2.2e-16 | 2.31 |
| **G and S subcentral** | 0.014 | 1.61 | 0.112 | 0.22 |
| **G and S transv frontopol** | -0.379 | -39.23 | <2.2e-16 | 5.29 |
| **G and S cingul-Ant** | -0.269 | -37.25 | <2.2e-16 | 5.02 |
| **G and S cingul-Mid-Ant** | -0.063 | -7.17 | 2.2e-9 | 0.97 |
| **G and S cingul-Mid-Post** | -0.048 | -5.38 | 1.6e-6 | 0.73 |
| **G cingul-Post-dorsal** | 0.098 | 12.06 | <2.2e-16 | 1.63 |
| **G cingul-Post-ventral** | -0.024 | -2.35 | 0.024 | 0.32 |
| **G cuneus** | -0.017 | -3.62 | 6.4e-4 | 0.49 |
| **G front inf-Opercular** | -0.006 | -0.75 | 0.454 | 0.10 |
| **G front inf-Orbital** | -0.009 | -0.58 | 0.564 | 0.08 |
| **G front inf-Triangul** | 0.136 | 14.10 | <2.2e-16 | 1.90 |
| **G front middle** | 0.120 | 13.78 | <2.2e-16 | 1.86 |
| **G front sup** | 0.085 | 15.89 | <2.2e-16 | 2.14 |
| **G Ins lg and S cent ins** | 0.045 | 2.90 | 0.005 | 0.39 |
| **G insular short** | -0.031 | -1.94 | 0.058 | 0.26 |
| **G occipital middle** | -0.022 | -1.70 | 0.094 | 0.23 |
| **G occipital sup** | -0.018 | -2.37 | 0.021 | 0.32 |
| **G oc-temp lat-fusifor** | -0.014 | -1.59 | 0.117 | 0.21 |
| **G oc-temp med-Lingual** | 0.078 | 13.08 | <2.2e-16 | 1.75 |
| **G oc-temp med-Parahip** | 0.086 | 8.28 | 3.4e-11 | 1.18 |
| **G orbital** | -0.134 | -33.59 | <2.2e-16 | 4.53 |
| **G pariet inf-Angular** | -0.043 | -3.89 | 2.8e-4 | 0.52 |
| **G pariet inf-Supramar** | -0.019 | -2.28 | 0.027 | 0.31 |
| **G parietal sup** | 0.172 | 20.29 | <2.2e-16 | 2.74 |
| **G postcentral** | 0.024 | 5.03 | 5.7e-6 | 0.68 |
| **G precentral** | -0.014 | -2.30 | 0.025 | 0.31 |
| **G precuneus** | 0.051 | 8.66 | 8.7e-12 | 1.17 |
| **G rectus** | 0.186 | 18.42 | <2.2e-16 | 2.48 |
| **G subcallosal** | 0.484 | 14.70 | <2.2e-16 | 1.98 |
| **G temp sup-G T transv** | 0.128 | 12.10 | <2.2e-16 | 1.63 |
| **G temp sup-Lateral** | 0.076 | 14.49 | <2.2e-16 | 1.95 |
| **G temp sup-Plan polar** | -0.195 | -14.01 | <2.2e-16 | 1.89 |
| **G temp sup-Plan tempo** | 0.089 | 8.28 | 3.4e-11 | 1.12 |
| **G temporal inf** | 0.024 | 2.95 | 0.005 | 0.40 |
| **G temporal middle** | -0.124 | -19.47 | <2.2e-16 | 2.62 |

| | | | | |
|---|---|---|---|---|
| **Lat Fis-ant-Horizont** | -0.282 | -14.99 | <2.2e-16 | 2.02 |
| **Lat Fis-ant-Vertical** | 0.204 | 10.79 | 4.3e-15 | 1.45 |
| **Lat Fis-post** | -0.291 | -54.74 | <2.2e-16 | 7.38 |
| **Pole occipital** | -0.483 | -65.71 | <2.2e-16 | 8.86 |
| **Pole temporal** | -0.116 | -25.05 | <2.2e-16 | 3.38 |
| **S calcarine** | 0.089 | 16.82 | <2.2e-16 | 2.27 |
| **S central** | 0.023 | 6.48 | 2.9e-8 | 0.87 |
| **S cingul-Marginalis** | -0.162 | -33.78 | <2.2e-16 | 4.56 |
| **S cincular insula ant** | -0.141 | -13.21 | <2.2e-16 | 1.78 |
| **S circular insula inf** | 0.185 | 29.87 | <2.2e-16 | 4.03 |
| **S circular insula sup** | 0.161 | 29.01 | <2.2e-16 | 3.91 |
| **S collat transv ant** | -0.067 | -5.59 | 7.6e-7 | 0.75 |
| **S collat transv post** | 0.060 | 3.20 | 0.002 | 0.43 |
| **S front inf** | 0.082 | 9.03 | 2.2e-12 | 1.22 |
| **S front middle** | -0.401 | -34.04 | <2.2e-16 | 4.59 |
| **S front sup** | 0.101 | 8.88 | 3.9e-12 | 1.20 |
| **S interm prim-Jensen** | -0.144 | -4.05 | 1.6e-4 | 0.55 |
| **S intrapariet and P trans** | -0.068 | -7.00 | 4.2e-9 | 0.94 |
| **S oc middle and Lunatus** | 0.120 | 9.90 | 9.6e-14 | 1.34 |
| **S oc sup and transversal** | -0.003 | -0.38 | 0.706 | 0.05 |
| **S occipital ant** | 0.049 | 2.06 | 0.044 | 0.28 |
| **S oc-temp lat** | -0.123 | -9.96 | 7.7e-14 | 1.34 |
| **S oc-temp med and Lingual** | 0.003 | 0.44 | 0.658 | 0.06 |
| **S orbital lateral** | -0.210 | -12.01 | <2.2e-16 | 1.62 |
| **S orbital med-olfact** | -0.081 | -9.73 | 1.7e-13 | 1.31 |
| **S orbital-H shaped** | 0.017 | 4.26 | 8.2e-5 | 0.57 |
| **S parieto occipital** | 0.028 | 8.77 | 5.7e-12 | 1.18 |
| **S pericallosal** | -0.073 | -5.40 | 1.5e-6 | 0.73 |
| **S postcentral** | 0.073 | 7.04 | 3.5e-9 | 0.95 |
| **S precentral-inf-part** | -0.012 | -1.64 | 0.106 | 0.22 |
| **S precentral-sup-part** | -0.031 | -3.99 | 2.0e-4 | 0.54 |
| **S suborbital** | 0.821 | 29.23 | <2.2e-16 | 3.94 |
| **S subparietal** | -0.198 | -20.84 | <2.2e-16 | 2.81 |
| **S temporal inf** | 0.210 | 17.92 | <2.2e-16 | 2.42 |
| **S temporal sup** | 0.019 | 2.49 | 0.016 | 0.34 |
| **S temporal transverse** | 0.189 | 11.73 | <2.2e-16 | 1.58 |

**Table S5:** Surface area lateralization and bias of manually labeled regions.

| Region | Metric | Mean | SD | *t* | *p* |
|---|---|---|---|---|---|
| **HG** | **original brains' lateralization** | 0.037 | 0.161 | 1.731 | 0.089 |
| | **flipped brains' lateralization** | -0.033 | 0.162 | -1.535 | 0.130 |
| | **bias** | 0.004 | 0.076 | 0.409 | 0.684 |
| **IFG pars opercularis** | **original brains' lateralization** | -0.002 | 0.146 | -0.090 | 0.929 |
| | **flipped brains' lateralization** | 0.003 | 0.128 | 0.175 | 0.862 |
| | **bias** | 0.001 | 0.077 | 0.122 | 0.903 |
| **IFG pars triangularis** | **original brains' lateralization** | 0.034 | 0.156 | 1.623 | 0.110 |
| | **flipped brains' lateralization** | -0.029 | 0.158 | -1.368 | 0.177 |
| | **bias** | 0.005 | 0.076 | 0.470 | 0.640 |
| **IFG pars orbitalis** | **original brains' lateralization** | -0.031 | 0.118 | -1.963 | 0.055 |
| | **flipped brains' lateralization** | 0.040 | 0.131 | 2.271 | 0.027 |
| | **bias** | 0.009 | 0.083 | 0.799 | 0.428 |

**Table S6:** Intraclass correlation coefficients of different parcellation schemes.

| | | transverse temporal (HG) | | IFG pars triangularis | | IFG pars orbitalis | | IFG pars opercularis | |
|---|---|---|---|---|---|---|---|---|---|
| | | ICC | *p* | ICC | *p* | ICC | *p* | ICC | *p* |
| **manual** | **real_lh** | 0.88 | 4.9e-14 | 0.85 | 1.1e-16 | 0.73 | 3.6e-10 | 0.76 | 5.2e-12 |
| | **real_rh** | 0.87 | 1.1e-16 | 0.83 | 1.3e-15 | 0.67 | 4.9e-7 | 0.71 | 2.7e-10 |
| ***aparc* default processing** | **real_lh** | 0.50 | 0.106 | 0.46 | 0.107 | 0.34 | 0.133 | 0.72 | 0.045 |
| | **real_rh** | 0.30 | 0.133 | 0.62 | 0.084 | 0.54 | 0.099 | 0.50 | 0.101 |
| **single atlas parcellation (lh atlas)** | **real_lh** | 0.93 | 6.5e-5 | 0.95 | 5.1e-4 | 0.94 | 3.3e-4 | 0.96 | 0.0028 |
| | **real_rh** | 0.94 | 0.000 | 0.96 | 7.9e-5 | 0.94 | 0.0018 | 0.96 | 0.003 |
| **single atlas parcellation (rh atlas)** | **real_lh** | 0.90 | 5.2e-4 | 0.96 | 0.0016 | 0.91 | 0.0056 | 0.96 | 8.9e-4 |
| | **real_rh** | 0.92 | 0.000 | 0.97 | 1.9e-4 | 0.93 | 0.0026 | 0.95 | 0.0019 |

**Table S7:** Systematic bias in surface area lateralization measurements obtained using symmetric registration but asymmetric atlas-based parcellation.

| Region | Mean Bias | *t* | *p* | Cohen's *d* |
|---|---|---|---|---|
| **bankssts** | 0.120 | 20.68 | <2.2e-16 | 2.788 |
| **caudal anterior cingulate** | 0.002 | 0.338 | 0.737 | 0.045 |
| **caudal middle frontal** | 0.020 | 7.573 | 4.8e-10 | 1.021 |
| **cuneus** | -0.022 | -6.182 | 8.694e-8 | 0.833 |
| **entorhinal** | -0.069 | -5.341 | 1.897e-6 | 0.720 |
| **frontal pole** | -0.191 | -20.373 | <2.2e-16 | 2.747 |
| **fusiform** | 0.043 | 13.866 | <2.2e-16 | 1.870 |
| **inferior parietal** | -0.123 | -35.268 | <2.2e-16 | 4.755 |
| **inferior temporal** | 0.077 | 30.358 | <2.2e-16 | 4.093 |
| **insula** | 0.003 | 1.653 | 0.104 | 0.223 |
| **isthmus cingulate** | 0.093 | 15.587 | <2.2e-16 | 0.093 |
| **lateral occipital** | 0.054 | 29.688 | <2.2e-16 | 4.003 |
| **lateral orbitofrontal** | 0.003 | 1.210 | 0.232 | 0.163 |
| **lingual** | -0.031 | -13.729 | <2.2e-16 | 1.851 |
| **medial orbitofrontal** | -0.100 | -26.942 | <2.2e-16 | 3.633 |
| **middle temporal** | -0.136 | -63.704 | <2.2e-16 | 8.590 |
| **paracentral** | -0.086 | -38.058 | <2.2e-16 | 5.132 |
| **parahippocampal** | 0.009 | 1.041 | 0.303 | 0.140 |
| **pars opercularis** | 0.183 | 46.790 | <2.2e-16 | 6.310 |
| **pars orbitalis** | -0.156 | -40.908 | <2.2e-16 | 5.516 |
| **pars triangularis** | -0.125 | -32.967 | <2.2e-16 | 4.445 |
| **pericalcarine** | -0.043 | -10.326 | 2.2e-14 | 1.392 |
| **postcentral** | -0.004 | -3.091 | 0.003 | 0.417 |
| **posterior cingulate** | 0.028 | 8.359 | 2.6e-11 | 1.127 |
| **precentral** | -0.003 | -2.007 | 0.050 | 0.271 |
| **precuneus** | -0.048 | -24.782 | <2.2e-16 | 3.342 |
| **rostral anterior cingulate** | 0.352 | 46.795 | <2.2e-16 | 6.310 |
| **rostral middle frontal** | -0.036 | -21.897 | <2.2e-16 | 2.953 |
| **superior frontal** | 0.035 | 28.163 | <2.2e-16 | 3.797 |
| **superior parietal** | 0.033 | 27.565 | <2.2e-16 | 3.717 |
| **superior temporal** | 0.031 | 15.035 | <2.2e-16 | 2.027 |
| **supramarginal** | 0.082 | 24.369 | <2.2e-16 | 3.286 |
| **temporal pole** | -0.055 | -6.507 | 2.6e-8 | 0.877 |
| **transverse temporal** | 0.300 | 57.538 | <2.2e-16 | 7.758 |

Statistics reflect one-sample t-tests against zero.

**Table S8:** Comparison of the systematic bias in surface area lateralization measurements obtained using left-atlas-parcellation and using default processing.

| Regions | bias when using left atlas only | bias in default processing | *t* | *p* |
|---|---|---|---|---|
| **bankssts** | 0.0045 | 0.188 | -18.890 | <2.2e-16 |
| **caudal anterior cingulate** | 0.0018 | -0.026 | 2.873 | 0.006 |
| **caudal middle frontal** | -0.0007 | 0.015 | -1.750 | 0.086 |
| **cuneus** | 0.0045 | 0.013 | -2.003 | 0.050 |
| **entorhinal** | 0.0067 | 0.109 | -6.872 | 6.6e-9 |
| **frontal pole** | 0.0112 | -0.152 | 18.865 | <2.2e-16 |
| **fusiform** | -0.0021 | 0.031 | -7.363 | 1.1e-9 |
| **inferior parietal** | 0.0047 | -0.090 | 15.116 | <2.2e-16 |
| **inferior temporal** | -0.0008 | 0.029 | -7.189 | 2.0e-9 |
| **insula** | 0.0049 | 0.020 | -1.781 | 0.081 |
| **isthmus cingulate** | 0.0150 | 0.059 | -6.043 | 1.4e-7 |
| **lateral occipital** | -0.0054 | 0.031 | -8.675 | 8.1e-12 |
| **lateral orbitofrontal** | 0.0033 | -0.027 | 6.205 | 8.0e-8 |
| **lingual** | -0.0011 | -0.036 | 11.232 | 9.6e-16 |
| **medial orbitofrontal** | 0.0119 | 0.027 | -2.119 | 0.039 |
| **middle temporal** | -0.0006 | -0.074 | 19.420 | <2.2e-16 |
| **paracentral** | 0.0016 | -0.119 | 28.927 | <2.2e-16 |
| **parahippocampal** | -0.0080 | -0.011 | 0.222 | 0.825 |
| **pars opercularis** | -0.0024 | 0.150 | -23.515 | <2.2e-16 |
| **pars orbitalis** | 0.0006 | -0.206 | 40.633 | <2.2e-16 |
| **pars triangularis** | -0.0037 | -0.173 | 39.504 | <2.2e-16 |
| **pericalcarine** | -0.0006 | -0.052 | 13.092 | <2.2e-16 |
| **postcentral** | 0.0023 | -0.020 | 6.478 | 2.9e-8 |
| **posterior cingulate** | 0.0179 | 0.037 | -3.637 | 0.001 |
| **precentral** | 0.0023 | 0.003 | -0.453 | 0.653 |
| **precuneus** | 0.0014 | -0.026 | 14.742 | <2.2e-16 |
| **rostral anterior cingulate** | 0.0074 | 0.216 | -15.703 | <2.2e-16 |
| **rostral middle frontal** | 0.0005 | -0.008 | 1.714 | 0.092 |
| **superior frontal** | 0.0013 | 0.044 | -10.297 | 2.4e-14 |
| **superior parietal** | -0.0013 | 0.032 | -8.054 | 8.1e-11 |
| **superior temporal** | 0.0044 | 0.050 | -18.497 | <2.2e-16 |
| **supramarginal** | -0.0015 | -0.007 | 0.592 | 0.556 |
| **temporal pole** | 0.0140 | 0.004 | 1.288 | 0.203 |
| **transverse temporal** | -0.0200 | 0.225 | -48.240 | <2.2e-16 |

**Table S9:** Comparison of the systematic bias in surface area lateralization measurements obtained using right-atlas-parcellation and using default processing.

| Regions | bias when using right atlas only | bias in default processing | *t* | *p* |
|---|---|---|---|---|
| **bankssts** | 0.0038 | 0.188 | -18.939 | <2.2e-16 |
| **caudal anterior cingulate** | 0.0075 | -0.026 | 3.426 | 0.001 |
| **caudal middle frontal** | -0.0002 | 0.015 | -1.721 | 0.091 |
| **cuneus** | 0.0026 | 0.013 | -2.314 | 0.024 |
| **entorhinal** | -0.0008 | 0.109 | -8.546 | 1.3e-11 |
| **frontal pole** | 0.0134 | -0.152 | 14.860 | <2.2e-16 |
| **fusiform** | 0.0004 | 0.031 | -6.742 | 1.1e-8 |
| **inferior parietal** | 0.0054 | -0.090 | 14.774 | <2.2e-16 |
| **inferior temporal** | -0.0006 | 0.029 | -7.563 | 5.0e-10 |
| **insula** | 0.0020 | 0.020 | -2.103 | 0.040 |
| **isthmus cingulate** | 0.0155 | 0.059 | -6.093 | 1.2e-7 |
| **lateral occipital** | -0.0048 | 0.031 | -8.771 | 5.7e-12 |
| **lateral orbitofrontal** | 0.0025 | -0.027 | 6.142 | 1.0e-7 |
| **lingual** | -0.0023 | -0.036 | 10.233 | 3.0e-14 |
| **medial orbitofrontal** | 0.0106 | 0.027 | -2.265 | 0.028 |
| **middle temporal** | -0.0025 | -0.074 | 19.156 | <2.2e-16 |
| **paracentral** | 0.0039 | -0.119 | 28.371 | <2.2e-16 |
| **parahippocampal** | -0.0100 | -0.011 | 0.071 | 0.943 |
| **pars opercularis** | -0.0017 | 0.150 | -25.230 | <2.2e-16 |
| **pars orbitalis** | -0.0030 | -0.206 | 41.270 | <2.2e-16 |
| **pars triangularis** | -0.0036 | -0.173 | 38.391 | <2.2e-16 |
| **pericalcarine** | 0.0000 | -0.052 | 14.535 | <2.2e-16 |
| **postcentral** | 0.0021 | -0.020 | 6.351 | 4.6e-8 |
| **posterior cingulate** | 0.0181 | 0.037 | -3.560 | 0.001 |
| **precentral** | 0.0019 | 0.003 | -0.590 | 0.557 |
| **precuneus** | 0.0018 | -0.026 | 12.601 | <2.2e-16 |
| **rostral anterior cingulate** | 0.0094 | 0.216 | -14.738 | <2.2e-16 |
| **rostral middle frontal** | 0.0028 | -0.008 | 2.298 | 0.025 |
| **superior frontal** | -0.0004 | 0.044 | -11.000 | 2.1e-15 |
| **superior parietal** | -0.0013 | 0.032 | -7.627 | 3.9e-10 |
| **superior temporal** | 0.0044 | 0.050 | -20.335 | <2.2e-16 |
| **supramarginal** | -0.0037 | -0.007 | 0.380 | 0.705 |
| **temporal pole** | 0.0174 | 0.004 | 1.530 | 0.132 |
| **transverse temporal** | -0.0225 | 0.225 | -38.751 | <2.2e-16 |

**Table S10:** Comparison of the systematic bias in surface area lateralization measurements obtained using left vs. right single-atlas-parcellation.

| Regions | bias when using left atlas only | bias when using right atlas only | *t* | *p* |
|---|---|---|---|---|
| **bankssts** | 0.0045 | 0.0038 | 0.164 | 0.870 |
| **caudal anterior cingulate** | 0.0018 | 0.0075 | -1.413 | 0.163 |
| **caudal middle frontal** | -0.0007 | -0.0002 | -0.279 | 0.781 |
| **cuneus** | 0.0045 | 0.0026 | 0.908 | 0.368 |
| **entorhinal** | 0.0067 | -0.0008 | 0.684 | 0.497 |
| **frontal pole** | 0.0112 | 0.0134 | -0.262 | 0.794 |
| **fusiform** | -0.0021 | 0.0004 | -1.623 | 0.110 |
| **inferior parietal** | 0.0047 | 0.0054 | -0.497 | 0.621 |
| **inferior temporal** | -0.0008 | -0.0006 | -0.104 | 0.918 |
| **insula** | 0.0049 | 0.0020 | 2.038 | 0.046 |
| **isthmus cingulate** | 0.0150 | 0.0155 | -0.159 | 0.874 |
| **lateral occipital** | -0.0054 | -0.0048 | -0.615 | 0.541 |
| **lateral orbitofrontal** | 0.0033 | 0.0025 | 0.567 | 0.573 |
| **lingual** | -0.0011 | -0.0023 | 0.982 | 0.331 |
| **medial orbitofrontal** | 0.0119 | 0.0106 | 0.563 | 0.576 |
| **middle temporal** | -0.0006 | -0.0025 | 1.120 | 0.268 |
| **paracentral** | 0.0016 | 0.0039 | -0.991 | 0.326 |
| **parahippocampal** | -0.0080 | -0.0100 | 0.412 | 0.682 |
| **pars opercularis** | -0.0024 | -0.0017 | -0.281 | 0.779 |
| **pars orbitalis** | 0.0006 | -0.0030 | 1.196 | 0.237 |
| **pars triangularis** | -0.0037 | -0.0036 | -0.033 | 0.974 |
| **pericalcarine** | -0.0006 | 0.0000 | -0.346 | 0.730 |
| **postcentral** | 0.0023 | 0.0021 | 0.156 | 0.877 |
| **posterior cingulate** | 0.0179 | 0.0181 | -0.068 | 0.946 |
| **precentral** | 0.0023 | 0.0019 | 0.391 | 0.697 |
| **precuneus** | 0.0014 | 0.0018 | -0.382 | 0.704 |
| **rostral anterior cingulate** | 0.0074 | 0.0094 | -0.276 | 0.784 |
| **rostral middle frontal** | 0.0005 | 0.0028 | -1.758 | 0.084 |
| **superior frontal** | 0.0013 | -0.0004 | 1.823 | 0.073 |
| **superior parietal** | -0.0013 | -0.0013 | 0.006 | 0.995 |
| **superior temporal** | 0.0044 | 0.0044 | 0.025 | 0.980 |
| **supramarginal** | -0.0015 | -0.0037 | 1.583 | 0.119 |
| **temporal pole** | 0.0140 | 0.0174 | -0.492 | 0.625 |
| **transverse temporal** | -0.0200 | -0.0225 | 0.635 | 0.528 |

**Table S11:** Surface area lateralization and bias of manually labeled regions with different parcellation schemes.

| Region | Metric | Manual | aparc default | single atlas parcellation (lh atlas) | single atlas parcellation (rh atlas) |
|---|---|---|---|---|---|
| HG | original brains' lateralization | 0.037 | 0.146 | 0.020 | 0.025 |
| | flipped brains' lateralization | -0.033 | 0.079 | -0.040 | -0.047 |
| | bias | 0.004 | 0.225 | -0.020 | -0.022 |
| IFG pars opercularis | original brains' lateralization | -0.002 | 0.080 | 0.009 | 0.002 |
| | flipped brains' lateralization | 0.003 | 0.070 | -0.011 | -0.003 |
| | bias | 0.001 | 0.150 | -0.002 | -0.001 |
| IFG pars triangularis | original brains' lateralization | 0.034 | -0.068 | 0.020 | 0.022 |
| | flipped brains' lateralization | -0.029 | -0.104 | -0.023 | -0.025 |
| | bias | 0.005 | -0.172 | -0.003 | -0.003 |
| IFG pars orbitalis | original brains' lateralization | -0.031 | -0.096 | 0.012 | 0.009 |
| | flipped brains' lateralization | 0.040 | -0.109 | -0.011 | -0.012 |
| | bias | 0.009 | -0.205 | 0.001 | -0.003 |

**Table S12:** Comparison of surface area lateralization obtained from manual labeling and from single-atlas parcellation.

| Region | Lateralization | Manual vs. lh_atlas | Manual vs. rh_atlas |
|---|---|---|---|
| **HG** | **original brains** | $t$ = -0.892, $p$ = 0.376 | $t$ = -0.647, $p$ = 0.520 |
| | **flipped brains** | $t$ = -0.335, $p$ = 0.739 | $t$ = -0.714, $p$ = 0.478 |
| **IFG pars opercularis** | **original brains** | $t$ = 0.586, $p$ = 0.560 | $t$ = 0.192, $p$ = 0.848 |
| | **flipped brains** | $t$ = -0.973, $p$ = 0.335 | $t$ = -0.053, $p$ = 0.957 |
| **IFG pars triangularis** | **original brains** | $t$ = -0.654, $p$ = 0.516 | $t$ = -0.573, $p$ = 0.569 |
| | **flipped brains** | $t$ = 0.266, $p$ = 0.791 | $t$ = 2.045, $p$ = 0.046 |
| **IFG pars orbitalis** | **original brains** | $t$ = 2.769, $p$ = 0.007 | $t$ = 2.634, $p$ = 0.011 |
| | **flipped brains** | $t$ = -3.030, $p$ = 0.004 | $t$ = -1.507, $p$ = 0.138 |
